# Independent representations of reward-predicting cues and reward history in frontal cortical neurons

**DOI:** 10.1101/2020.01.23.916460

**Authors:** Masashi Kondo, Masanori Matsuzaki

## Abstract

The transformation of sensory inputs to appropriate goal-directed actions requires estimation of sensory-cue values based on outcome history. To clarify how cortical neurons represent an outcome-predicting cue and actual outcome, we conducted wide-field and two-photon calcium imaging of the mouse neocortex during performance of a classical conditioning task with two cues with different water-reward probabilities. Although licking behavior dominated the area-averaged activity over the whole dorsal neocortex, dorsomedial frontal cortex (dmFrC) affected other dorsal frontal cortical activities, and its inhibition extinguished differences in anticipatory licking between the cues. In individual frontal cortical neurons, the reward-predicting cue was not simultaneously represented with the current or past reward, but licking behavior was frequently multiplexed with the reward-predicting cue and current or past reward. Deep-layer neurons in dmFrC most strongly represented the reward-predicting cue and recent reward history. Our results suggest that these neurons ignite the cortical processes required to select appropriate actions.

## Introduction

The transformation of sensory signals to action is crucial for survival in all animals. An outcome following a sensory signal updates the value of the sensory signal. This updated value is then used in the next response to the same sensory signal to evoke a specific action to obtain or avoid an outcome. In decision-making tasks that require goal-directed actions to successfully receive a reward, task-related and movement-related activities are widely distributed over the whole dorsal neocortex and whole brain (Allen et al., 2017; Musall et al., 2019; Pinto et al., 2019; Steinmetz et al., 2019).

In the rodent cerebral cortex, the secondary motor area (M2/MOs), which is a broad anatomically defined area (Allen Brain Atlas or Barthas and Kwan, 2017; Ebbesen et al., 2018; Franklin and Paxinos, 2007), plays critical roles in value estimation, action selection, working memory, and motor planning (Allen et al., 2017; Gilad et al., 2018; Guo et al., 2017; Murakami et al., 2014; Pinto et al., 2019; Siniscalchi et al., 2016; Sul et al., 2011; Svoboda and Li, 2018). In addition, the activation of lateral and medial areas of M2 induces tongue and forelimb movements, respectively, and orofacial movements are also related to M2 (Ebbesen et al., 2018; Guo et al., 2017; Hira et al., 2015; Komiyama et al., 2010; Mimica et al., 2018; Tennant et al., 2011; Wang et al., 2017; Yoshida et al., 2009). The type of action required to obtain a reward varies across different decision-making studies, and the representation of each M2 neuron is frequently assigned to one type of task-related information, that which is dominantly represented (Musall et al., 2019; Stringer et al., 2019).

To clarify action-type-independent decision-making-related information in M2, it is necessary to measure the activities of individual neurons within subdivisions of M2 during simple tasks and to analyze how individual neurons encode the multiple types of information. We assumed that the neuronal representations of the cue value and outcome information in classical conditioning, a condition that does not require the learning of an association between a specific action and its consequence, is the basis for the processes of the transformation of sensory cues to appropriate motor actions. In classical conditioning tasks, striatal neurons represent information on reward-predicting cues (cue values), present reward (including reward prediction error), and recent reward history (reward in the preceding trial), as well as behaviors such as licking (Bloem et al., 2017; Yoshizawa et al., 2018). However, it is still unclear how the dorsal neocortex, including M2, represents such information, which is generally required for decision-making.

To extract these information types, we trained head-fixed mice to perform a classical conditioning task with two sound cues assigned to different probabilities of water delivery (Bloem et al., 2017; Oyama et al., 2015; Shin et al., 2018). We conducted wide-field calcium imaging of the entire dorsal cortex (Allen et al., 2017; Makino et al., 2017; Musall et al., 2019) during the training sessions. In addition, in the late stage of learning, we conducted two-photon calcium imaging of three dorsal frontal areas up to a depth of 800 μm from the cortical surface (dorsomedial frontal cortex [dmFrC] corresponding to antero-medial M2, dorsolateral frontal cortex [dlFrC] corresponding to antero-lateral M2 [or occasionally referred to as the anterolateral motor area; ALM], and the primary motor cortex [M1] corresponding to the caudal forelimb motor area (Franklin and Paxinos, 2007; Hira et al., 2013a, 2013b; Komiyama et al., 2010; Svoboda and Li, 2018; Tennant et al., 2011), and the medial prefrontal cortex at depths of 800–1200 μm (mPFC; roughly corresponding to the dorsal part of the prelimbic area). We found that in all dorsal cortical areas, licking-related activity dominated throughout the sessions, while information regarding the reward-predicting cue emerged over the dorsal cortex and information regarding the recent reward history emerged in the frontal cortex during learning. Individual dmFrC neurons strongly but separately represented these types of task-related information. Our results suggest that dmFrC can integrate different types of task-related information to select the appropriate action in the decision-making process.

## Results

### Classical conditioning task with two sound cues assigned to different reward probabilities

We trained head-fixed mice to perform a classical conditioning task where two different tones acting as conditioned stimuli (CS, cues A and B) were assigned to different probabilities of the delivery of an unconditioned stimulus (US), which was a reward of a drop of water (P = 0.7 for cue A and 0.3 for cue B; Figure 1A; Bloem et al., 2017). Cue A or B was randomly presented in each trial. The cue was presented for 1.5 s (cue period) followed by a delay period of 0.5 s, and then a drop of water was delivered near to the mouth from a spout, according to the determined probability. The head-fixed mice were able to lick freely.

**Figure 1.**
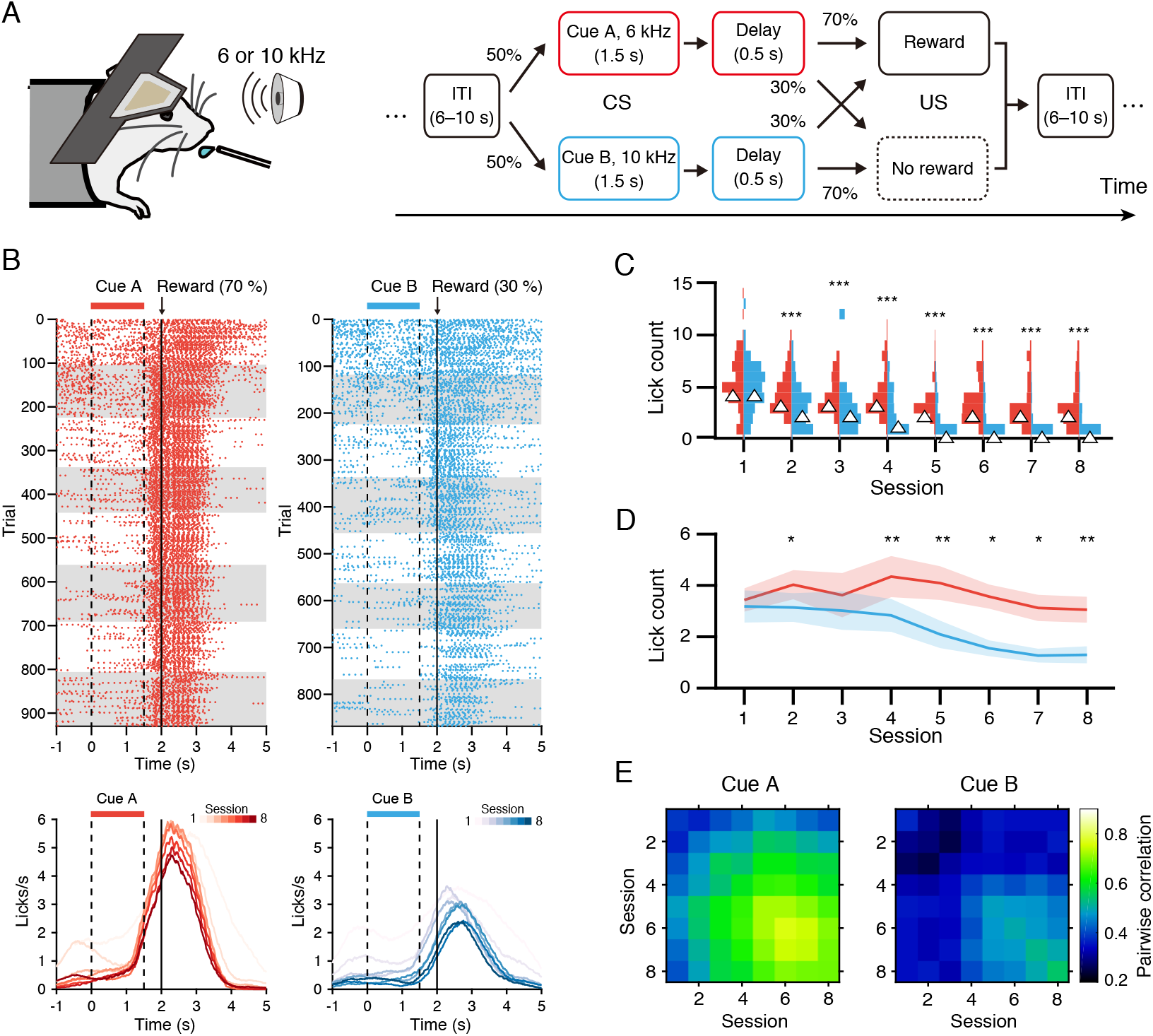
Mice showed anticipatory behavior through learning. (**A**) Schematic illustration of a classical conditioning task with different reward probabilities. When a trial started, either tone cue A or tone cue B (CS) was randomly presented to head-fixed mice for 1.5 s, and then after a 0.5 s delay, a drop of water (US) was delivered according to a probability of 0.7 (cue A) or 0.3 (cue B). (**B**) Changes in licking behavior for each CS (left, cue A; right, cue B) trial over eight sessions in one mouse. Gray shaded areas indicate even numbers of sessions. The lower panel shows the time course of the average lick-rate in each session. SEM is not shown to facilitate clear data visualization. (**C**) Changes in the histogram of the count of licks during 2 s after the CS onset (anticipatory licks) in the same mouse as in (**B**). Red, cue A. Blue, cue B. Triangles indicate the median for each cue. ***: P < 0.001, Wilcoxon rank-sum test. (**D**) Changes in the count of the anticipatory licks for all animals (n = 6). Shaded areas are SEMs. *: P < 0.05, **: P < 0.01, Wilcoxon signed-rank test. (**E**) Intra- and inter-session trial-by-trial correlations of the anticipatory licking. The correlation coefficients were averaged over all animals.

At the beginning of training, which followed 2–3 pretraining sessions in which the reward was delivered at 100% after both cue presentations, the number of licks during a 2 s period after the cue onset (anticipatory licking) was similar between trials with cue A (cue A trials) and cue B presentations (cue B trials; Figures 1B–1D). However, the number of anticipatory licks in cue B trials became lower than the number in cue A trials as the training progressed (Figures 1B–1D). Intra- and inter-session trial-by-trial correlations of the anticipatory lick-rate in cue A trials reached a plateau around session 6 (Figure 1E, left). By contrast, the increase in the correlation in cue B trials was smaller and slower (Figure 1E, right). These results indicate that the mice learned that cues A and B were associated with different reward probabilities, and the anticipatory licking behavior in cue A trials therefore became robust and stable.

### Cortex-wide neuronal activity changes in response to the learning of associations between the sensory cue and reward

To examine how neuronal activity in the dorsal cerebral cortex changed during learning of the task, we conducted transcranial wide-field calcium imaging of the entire dorsal cortex in transgenic mice (Figures 2A–2D) over eight daily sessions. These mice widely expressed a red genetically-encoded calcium indicator (R-CaMP1.07) in cortical excitatory neurons (Emx1-IRES-Cre::CaMKII-tTA::TITL-R-CaMP1.07; Bethge et al., 2017). Although there is some abnormal activity in the frontal cortex of transgenic mice in which GCaMPб is widely expressed in the whole brain (e.g., Emx1-Cre, Steinmetz et al., 2017), apparent abnormal activity was not observed in any dorsal cortical area of the transgenic mice that we used (Figure S1A). Twelve dorsal cortical areas were set as regions of interest (ROIs) according to the Allen Common Coordinate Framework version 3 (Figure 2E), with these including the dmFrC and dlFrC areas in M2.

**Figure 2.**
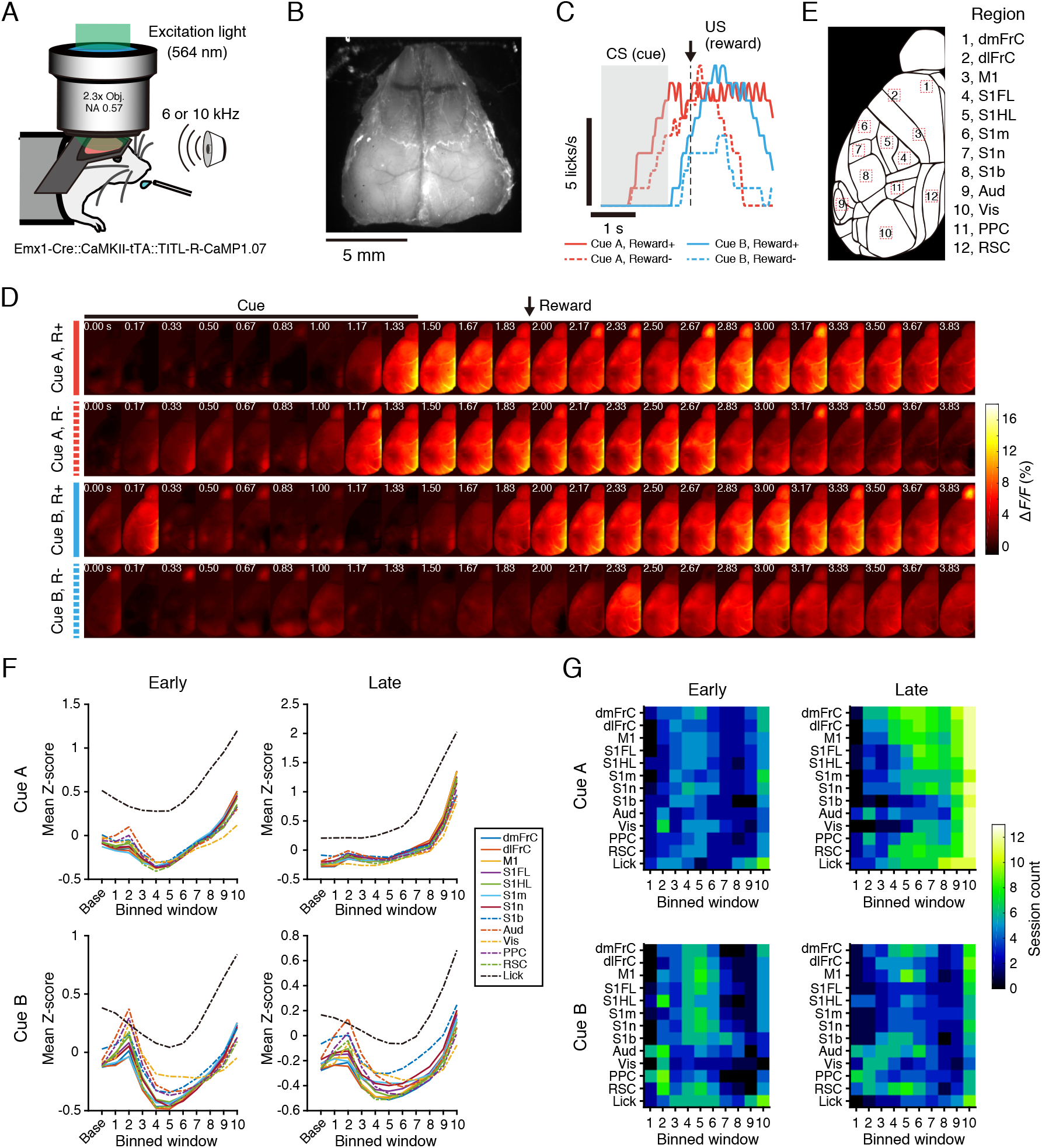
Wide-field calcium imaging revealed cortex-wide neuronal activities over the task sessions. (**A**) Schematic illustration of wide-field calcium imaging in Emx1-Cre::CaMKII-tTA::TITL-R-CaMP1.07 mice during task performance. (**B**) A representative transcranial fluorescence image. (C) Examples of the lick-rate in four trials: cue A with reward delivery (red solid line), cue A without reward delivery (red dashed line), cue B with reward delivery (blue solid line), and cue B without reward delivery (blue dashed line). (**D**) Montage images of wide-field neuronal activities in the left hemisphere in the four trials shown in (C). (**E**) Locations of the twelve ROIs superimposed on the top view of the Allen common coordinate framework ver.3. dmFrC, dorsomedial frontal cortex; dlFrC, dorsolateral frontal cortex; M1, primary motor cortex; S1FL, primary forelimb sensory cortex; S1Hl, primary hindlimb sensory cortex; S1m, primary mouth sensory cortex; S1n, primary nose sensory cortex; S1b, primary barrel sensory cortex; Aud, auditory cortex; Vis, visual cortex; PPC, posterior parietal cortex; RSC, retrosplenial cortex. (**F**) Neuronal activities in the twelve ROIs and the lick-rate in early (sessions 1 and 2, left) and late (sessions 7 and 8, right) stages of learning. The neuronal activities and lick-rates during the 2 s period after the cue onset were averaged for each 200 ms bin, and the baseline activity (base) and lick-rate were defined as the average over the 500 ms period before the cue onset. (**G**) The number of sessions with significant differences (P < 0.05, FDR corrected) in lick-rate or neuronal activity in each ROI between the baseline window and each time-bin (12 sessions = 6 mice × 2 sessions in each stage; see Methods for details).

First, we examined how the trial-averaged activity in each of the ten 200-ms bins during the cue and delay periods (total, 2 s) differed from the baseline activity (Figures 2F and 2G). In cue A trials, the auditory area showed increased activity in the second time-bin in both early (sessions 1–2) and late stages (sessions 7–8) of learning. In the late stage of learning, the activity in the frontal areas including dmFrC, dlFrC, and M1, as well as the lick-rate, increased after the second or third time-bin, and many other areas showed increased activity after the fifth bin. In cue B trials, the posterior cortex including the auditory area and posterior parietal cortex (PPC) showed increased activity in the first or second bin in both stages of learning. The latter tendency was observed even in the pretraining sessions, and the activity in the posterior cortex was larger in cue B trials than in cue A trials (Figures S1B and S1C). These results indicate that the rapid rise of the dorsal frontal cortex activity in cue A trials in the late stage of learning reflected learning of the reward-predicting cue, rather than simply reflecting the sensation of the cue A tone.

### Cortex-wide neuronal activity and licking are highly synchronized, but the cortical information flow is inferred to emerge from dmFrC

Next, we examined whether the neuronal activity during the cue and delay periods was correlated with the dorsal cortical areas. In each trial type, the cross-correlation was maximized at zero-lag between any pair of areas (Figure 3A). Additionally, trial-by-trial linear correlations in activity were high in many pairs in both stages of learning, especially the correlations between the dorsal frontal cortical areas (dmFrC, dlFrC, M1, primary sensory forelimb [S1FL], and hindlimb areas [S1HL]) (Figure 3B).

**Figure 3.**
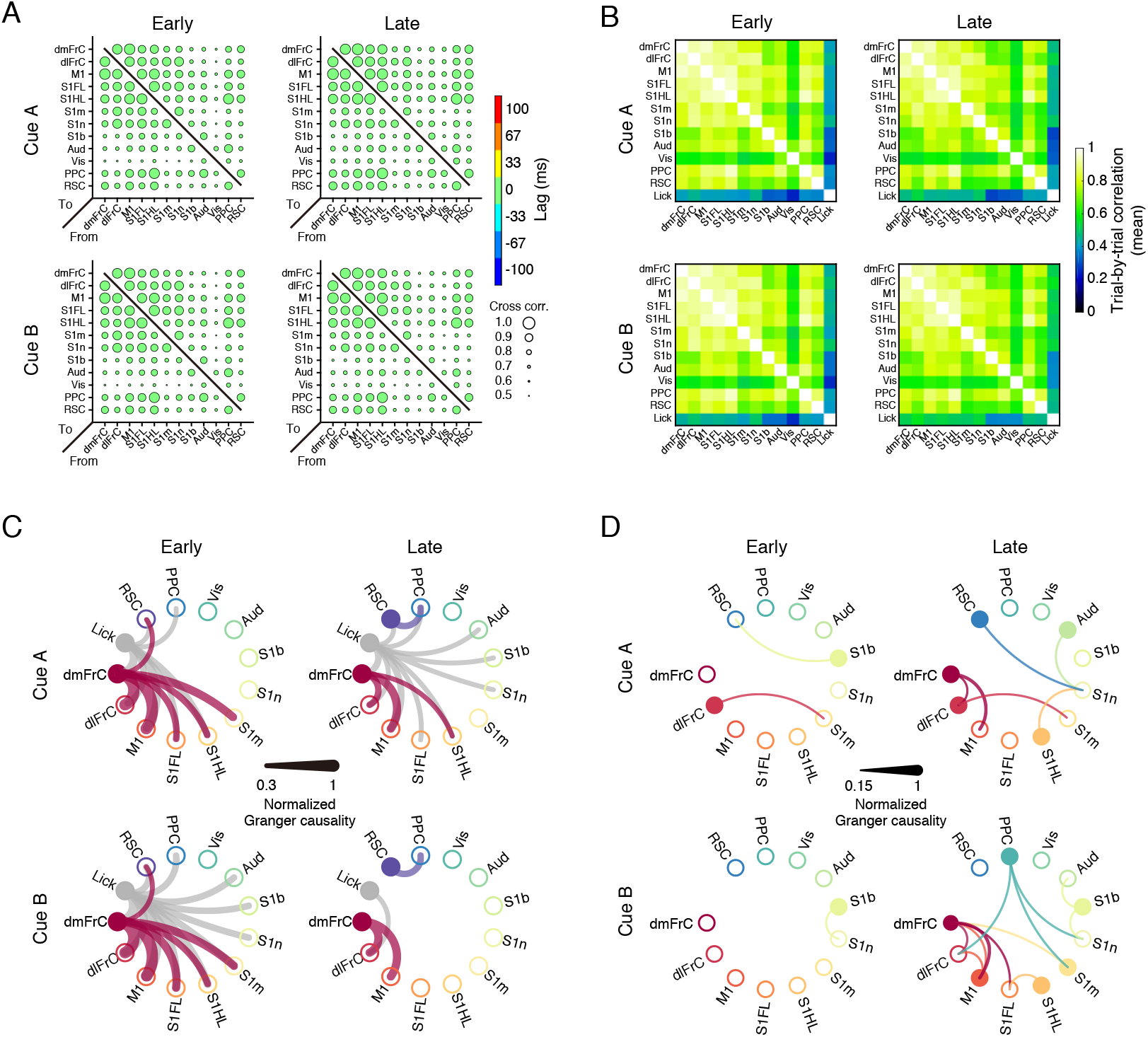
Area-averaged activities and licking behavior were highly synchronized, but an information flow unrelated to licking behavior emerged within the frontal cortices. (**A**) Cross-correlations in activity during the 2 s period after the cue onset between the areas along the horizontal and vertical axes. The color of circle indicates the time lag at which the peak correlation coefficient occurred, and all were assigned to a lag time of zero. The size of the circles indicates the peak amplitude of the correlation coefficient. The values for each stage are averaged over 12 sessions. (**B**) Trial-by-trial pairwise linear correlations between neuronal activities of different regions and between each neuronal activity and lick-rate. The data are for the 2 s period after the cue onset. (**C**) Granger causality between neuronal activities of different regions and between neuronal activity and lick-rate. If the causality was above 0.3, the causal root and sink are connected with a line. The color of the line is the same as the color assigned to the area of the causal root, with the circle of the causal root being filled. The thickness of the line corresponds to the strength of the causality. (**D**) Granger causality between neuronal activities obtained in only those trials without any anticipatory licking. If the causality was above 0.15, the causal root and sink are connected with a line.

By contrast, pairwise conditional Granger causality analysis (Barnett and Seth, 2014; Makino et al., 2017) suggested that dmFrC was the critical area for predicting the activity of dlFrC and M1 in both cue A and cue B trials (Figures 3C, S2A, and S2B), and that the anticipatory lick-rate predicted the activity in many areas in cue A trials in the early and late stages, and in cue B trials in the early stage. Therefore, licking might have dominantly affected the activity of entire dorsal cortices, while the dmFrC activity might have affected the putative downstream areas independent of the licking events. To test this, we analyzed only those trials without anticipatory licking. Although the interareal relations with the Granger causality were very weak, dmFrC still predicted dlFrC and M1 activities in both cue trials in the late stage of learning (Figure 3D). This suggests that the learning induced strong associations between dmFrC and dlFrC, and between dmFrC and M1 (Figure 3D), although the licking behavior affected many cortical areas throughout learning.

We also examined the changes in cortex-wide population activity through learning. We performed factor analysis (Harvey et al., 2012; Makino et al., 2017) to decompose the neural activities of 12 cortical regions into five factors representing the population activity. As the learning stage progressed, the inter-trial variability of the population activity in cue A trials decreased, and the distance in the neuronal trajectories between cue A and cue B trials increased (Figures S2C-S2E). These results might suggest that the cortex-wide neuronal activity in cue A trials converged to reduce the trial-by-trial variability through learning of the high reward outcomes.

However, the effect of learning on the convergence of the cortex-wide activity could also be explained by the licking behavior because its trial-by-trial variability also decreased as the learning progressed (Figure 1E), and such movements can have a large effect on the neuronal activity as a whole (Musall et al., 2019; Steinmetz et al., 2019; Stringer et al., 2019). Therefore, we compared the trial-by-trial correlation of the lick-rate and the trial-by-trial correlation of the neuronal trajectories in each learning stage and CS presentation, and found that these correlations showed a strong association (Figure S2F). The association extent was similar between the early and late stages of learning, and the trial-by-trial correlation in the cortex-wide activity was not high when the trial-by-trial correlation of lick-rate was low. These results suggest that the cortex-wide population activity was strongly related to the licking behavior throughout learning.

### Dorsomedial frontal cortex is necessary for value-dependent anticipatory licking

Despite the large effect of the licking behavior on the activities of the dorsal cortical areas, the information flow (or neuronal activation around the dorsal cortex) appeared to be initiated from the frontal cortex, especially the dmFrC (Figure 3D). To test the relationship between the frontal cortical activity and anticipatory licking behavior in the current conditioning task, we photoinhibited three frontal areas (dmFrC, dlFrC, and M1) and the primary somatosensory cortex barrel region (S1b; as an assumed negative control) in VGAT-ChR2-EYFP mice (Zhao et al., 2011) during 3 s periods after the cue onset (Figures 4A and 4B).

**Figure 4.**
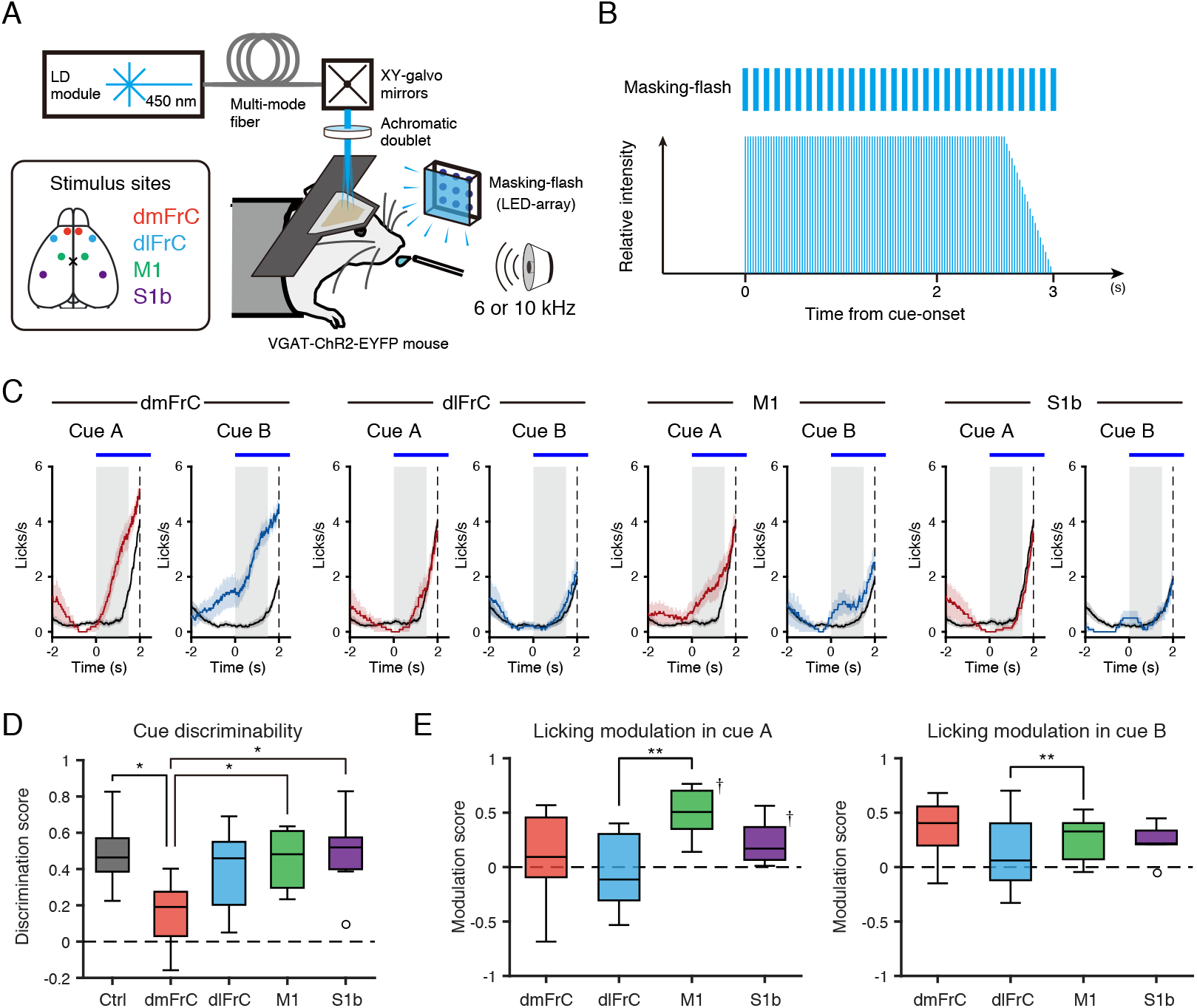
Photoinhibition of the dorsomedial frontal cortex abolished the difference in anticipatory licking between the cues. (**A, B**) A schematic illustration of a photoinhibition experiment with VGAT-ChR2-EYFP mice. Photoinhibition light was masked by flashes of light. Inset, stimulated locations. One of four cortical areas (dmFrC, dlFrC, M1, and S1b) was bilaterally photoinhibited in randomly chosen half of the trials. Flashing to mask photostimulation was performed in all trials. B, The intensity and timing of photostimulation during the task. (**C**) Examples of the trial-averaged time course of the lick-rate for each photoinhibition location and sound cue. Lick-rates are shown in mean ± SEM. Left: cue A, right: cue B. The red and blue lines are lick-rates when the photoinhibition was performed, and the black lines are lick-rates when it was not performed. Gray shading indicates the cue presentation period. The blue horizontal bars at the top of the plot indicate the photoinhibition period. (**D**) Photoinhibition-induced changes in cue discriminability. The cue discriminability was estimated by the discrimination score (see Methods). The box edges indicate the first and third quartiles; the inner line is the median over eight mice; whiskers represent minimum and maximum values except for outliers; the circle represents an outlier outside 1.5 times the IQR. *: P < 0.05, Friedman test with multiple correction by dunn-sidak method. (**E**) Photoinhibition-induced modulation of the lick-rate for each cue. The modulation was estimated by the modulation score (see Methods). Left: cue A, right: cue B. **: P < 0.01, Friedman test with multiple correction by dunn-sidak method. †: P < 0.05 vs zero, sign test, FDR corrected.

In the pretraining sessions, only photoinhibition of dlFrC decreased the lick-rate during the cue and delay periods (Figure S3). By contrast, when we conducted the photoinhibition after the training was completed (see Methods), only photoinhibition of dmFrC decreased the discriminability in the anticipatory licking between cue A and cue B (Figures 4C–4E). This suggests that the association with the cue-type dependent anticipatory licking after learning was stronger in dmFrC than in the other frontal cortical areas.

### Licking-, cue-, and reward-related activities of individual neurons in the dorsal frontal cortex and mPFC

It is not possible to detect the activity of individual neurons or neuronal activity in deep layers on transcranial wide-field calcium imaging. In addition, the fluorescent changes in each area also include the activity of axons from other brain areas (Allen et al., 2017). Therefore, we conducted two-photon calcium imaging to detect the activities of multiple neurons in both superficial and deep layers at a single-cell resolution. The wide-field imaging and photoinactivation experiments indicated that dorsal frontal cortex activity would be strongly related to cortex-wide activity and anticipatory licking during the cue and delay periods. Therefore, we injected an adeno-associated virus (AAV) carrying jRGECO1a (Dana et al., 2016) into dmFrC, dlFrC, and M1. In addition, after injecting the AAV into the prelimbic area, we conducted two-photon calcium imaging of the mPFC underneath the dmFrC, a region that cannot be detected with wide-field imaging (Kondo et al., 2017).

We trained the mice expressing jRGECO1a for more than eight sessions, and then conducted two-photon imaging of the same areas through the cranial window (Figures 5A–5C). From a total of 19 mice, we analyzed data from 5680 dmPFC neurons (53 imaging fields), 6232 dlFrC neurons (50 imaging fields), 3073 M1 neurons (41 imaging fields), and 2032 mPFC neurons (16 imaging fields). First, we defined the imaged neurons that showed activity that correlated with lick-rate (Pearson’s correlation coefficient of > 0.3 with statistical significance [P < 0.05]) as licking-related neurons (Figure 5D). As expected from the results of the wide-field imaging, more than 39% of the neurons at all depths in three dorsal cortical areas were classified into licking-related neurons (Figure 5E). By contrast, the proportion of the licking-related neurons in the mPFC was smaller at approximately 20%.

**Figure 5.**
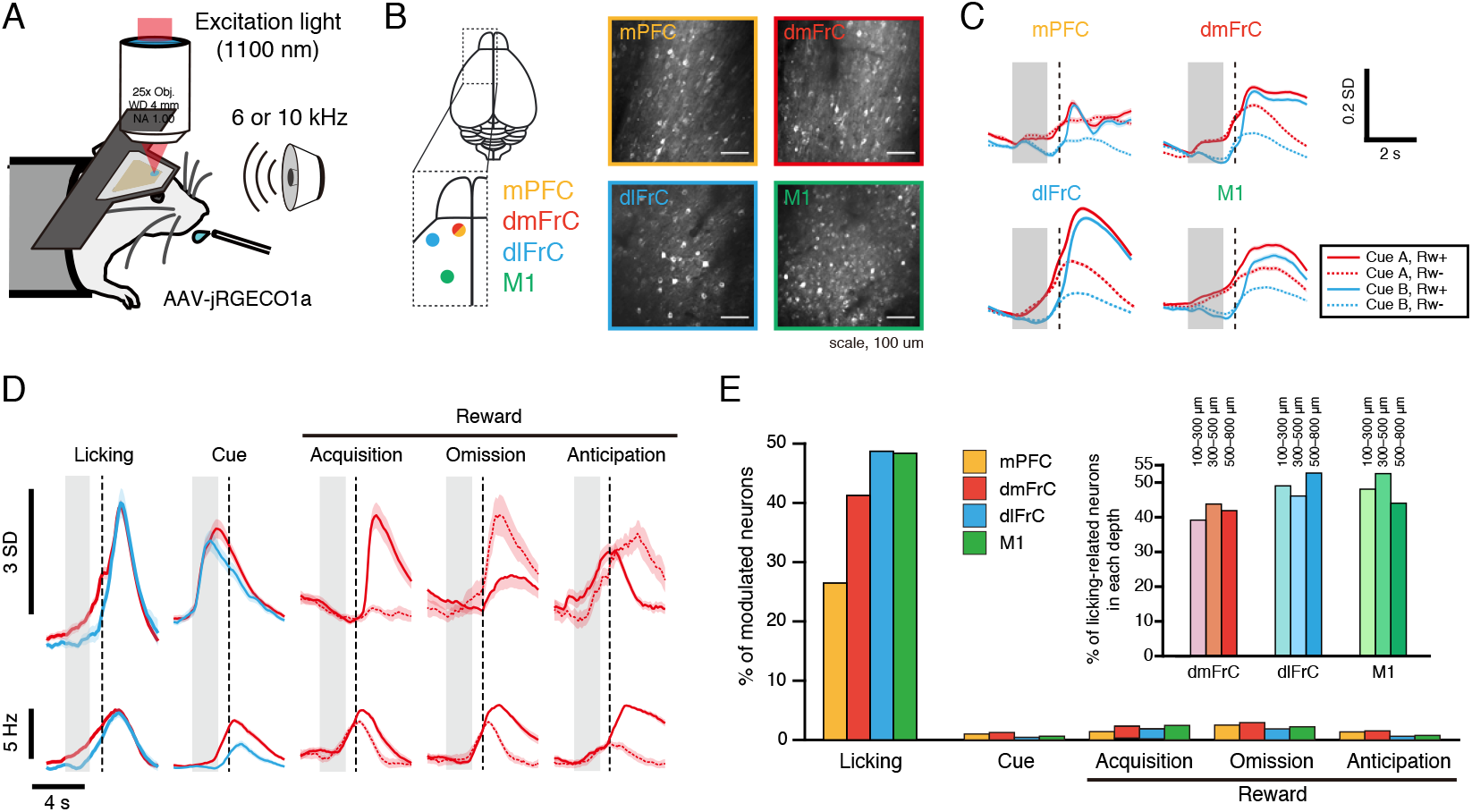
Licking behavior dominated a large portion of the single-cell activity in the frontal cortices. (**A**) A schematic diagram of two-photon calcium imaging in head-fixed mice performing the task. (**B**) Left, four imaged fields: mPFC (yellow), dmFrC (red), dlFrC (cyan), and Ml (green). mPFC and dmFrC were located at the same AP-ML coordinates and differed in depth from the cortical surface (dmFrC: 100–800 μm, mPFC: > 800 μm). Right, representative time-averaged fluorescent images in each field. (**C**) Population activities of each frontal cortical area. For each area, trial-type-averaged neuronal activities were averaged across neurons from all depths. Red and blue lines correspond to activities in cue A and cue B trials, respectively. Solid and dashed lines correspond to trials with and without reward delivery, respectively. (**D**) Examples of the activity of five neurons with task-related activity defined according to canonical criteria (see Methods for details). Top, trial-averaged neuronal activity in different trial conditions. Bottom, corresponding lick-rate. From left to right panel: licking-related, cue (red and blue, trials with cue A and B, respectively), reward acquisition, reward omission, and reward anticipation neurons (solid lines: trials with reward delivery, dashed lines: trials without reward delivery). (**E**) The ratios of the five types of neurons defined by the canonical criteria in each area. Yellow: mPFC, red: dmFrC, blue: dlFrC, green: M1. Inset, the proportion of the licking-related neurons at each depth (100–300, 300–500 or 500–800 μm from the cortical surface) in dmFrC, dlFrC, and M1.

Next, neurons other than these licking-related neurons were classified into four types according to whether activity was related to the cue, reward acquisition, reward omission, or reward anticipation on the basis of differences in the average activity between specific task-related time-bins (Wagner et al., 2017; Figures 5C–5E; see Methods for details). However, in this simple classification, only approximately 10% of the neurons in each area were cue- and outcome-related.

### Multiple linear regression separately extracts the task-related and licking-related activities of individual neurons

To finely extract the cue and outcome information from individual neurons, irrespective of whether licking information was present or not, we next applied multiple linear regression to the activity of each neuron (Allen et al., 2017; Chen et al., 2017b; Siniscalchi et al., 2019) for each time-frame of 100 ms during a 10 s trial period (from 2 s before to 8 s after the onset of the cue presentation; 101 frames in total), using predictor variables of cue (cue A or cue B), outcome (presence or absence of the reward), and licking (Figures 6A, S4 and S5A; see Methods). In addition to the cue and outcome information in the current trial, we used the information regarding the cues and outcomes in one and two trials before the current trial as predictor variables (cue history and reward history). The Z-scored lick-rates in the current trial were averaged every 2 s; therefore, three cue variables, three outcome variables, and five licking variables were used as predictor variables in each trial (Figure S4A). The population-level prediction accuracy was highest in dmFrC and lowest in M1 (Figure S4B). Within dmFrC, the layer at depths of 500–800 μm had higher prediction accuracy than other depths (Figure S4B).

**Figure 6.**
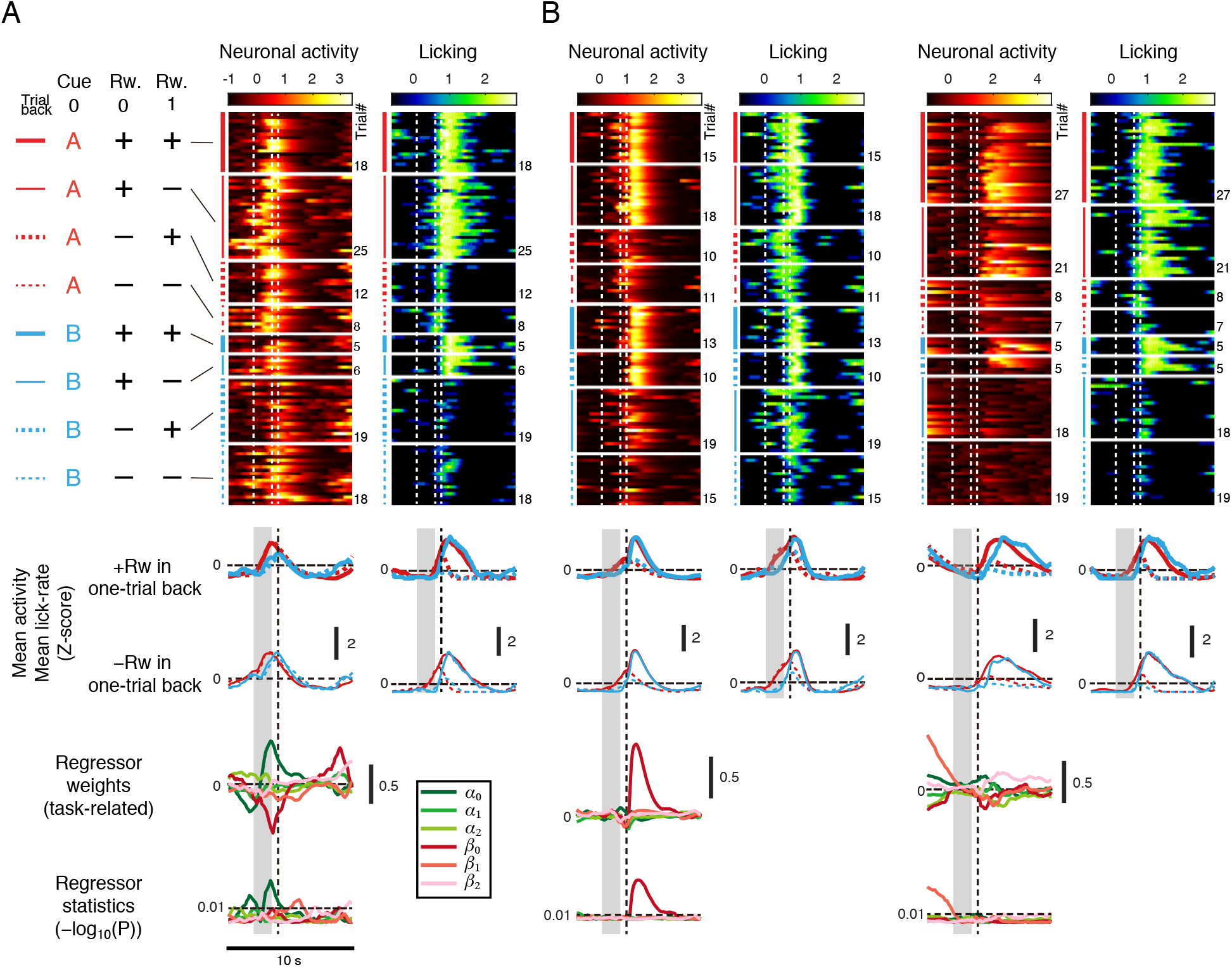
Encoding model extracted dynamics of cue and outcome variables from licking-related neurons. Three representative examples of neuronal activity, task-related regressor weights and statistics, and licking behavior. Red and blue in the mean ΔF/F traces correspond to cue A or cue B trials, respectively. Solid and dashed lines correspond to trials with or without the current reward delivery, respectively. Thick and thin lines correspond to trials with or without one-trial back reward delivery, respectively. In the weights and P-values of the regressor, greenlike colors correspond to the cue variables and red-like colors to the outcome variables. The color depths correspond to zero-, one-, or two-trials-back from dark to pale color. In this figure, all regressor weights are shown as original estimates, regardless of their P-values. The P-values of the regressor weights are shown using a —log_10_ scale. For the mean neuronal activity and lick-rate, SEM was not shown to facilitate clear data visualization.

The regression allowed extraction of the weights of the cue and outcome variables from the licking-related neurons and other neurons (Figures 6 and S5). To examine the types of task-related information possessed by the neurons in each area, we estimated the proportions of neurons that had predictor variables with P-values of < 0.01 at each time-frame at the three depth ranges of 100–300, 300–500, and 500–800 μm from the cortical surface in dmFrC, dlFrC, and M1, and at a depth of 800–1200 μm in mPFC (Figure 7A). At all depths in dmFrC, the current cue was a predictor variable in approximately 10% of the neurons at the peak time point around the delay period. In the deep layer (300–800 μm) of the dlFrC, ~5% of neurons had the current cue as a predictor variable at the peak time point during the delay period (Figure 7A). In all areas at all depths, 5–15% of neurons had substantial weights for the current outcome variable at the peak time point after the reward delivery (by binomial test). In addition, 10–20% of neurons in mPFC, dmFrC, and dlFrC had substantial weights for the one-trial-back outcome variable (recent reward history) at the peak time point during the pre-cue period (from 2 s to 0 s before cue onset). The weights of the cues for one and two-trials-back and the outcome of the two-trials-back did not exceed the chance level at any time point. The proportions of neurons whose activities were explained by the licking variables were higher than the chance level in all areas (Figure S4C).

**Figure 7.**
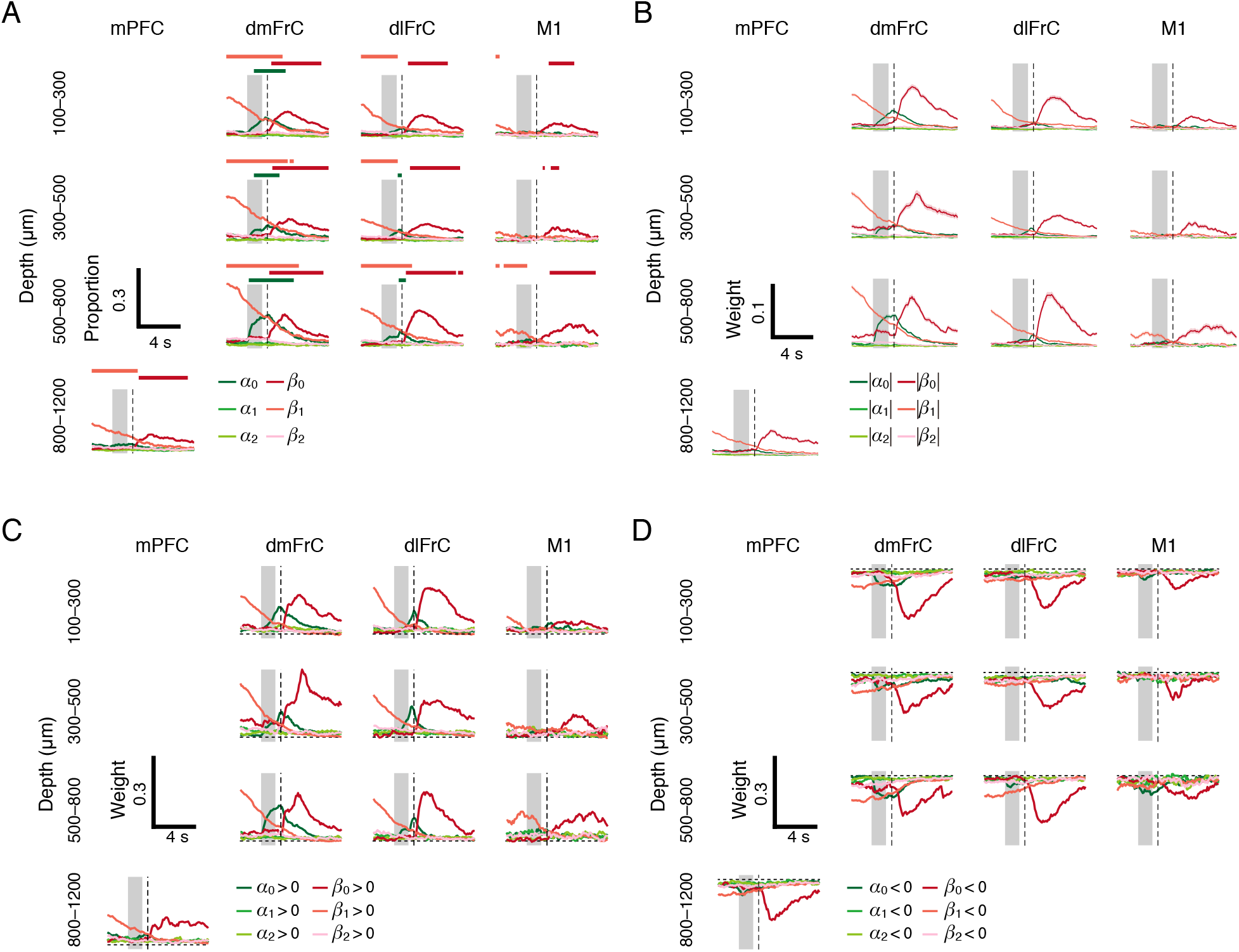
Time course of regressor weights at each depth in each area. (**A**) The proportion of neurons with regressors with large weights (P < 0.01) for each variable in each field and each depth. The upper horizontal lines indicate that the cell population had significant corresponding regressors above the chance level (binomial test, P < 0.01) at that time point. (**B**) Mean absolute values of the regressor weights for each field and each depth. Only weights with a P value of < 0.01 at each time point in each neuron were used in the calculation. SEM is not shown to facilitate clear data visualization. (**C, D**) Mean positive values (C) or negative values (D) of the regressor weights at each cortical depth in each field. Only neurons that had either positive (**C**) or negative (D) peak magnitude weights with P-values of < 0.01 were used in the calculation. SEM is not shown to facilitate clear data visualization.

Similar results were obtained when the absolute value of the weight of each variable was estimated (Figure 7B), and the results were also similar when neurons with positive or negative weights for each variable were separately grouped (Figures 7C and 7D). A prominent result was that dmFrC had more neurons with negative weights for the current cue (that is, higher activity in cue B trials than in cue A trials) than in the other areas. These results indicate that the reward-predicting cue was strongly represented by dmFrC neurons, the recent reward history was represented by dmFrC, dlFrC, and mPFC neurons, and the current outcome and lick-rate were represented by neurons in all four areas.

The task-related variables (three cue and outcome variables) may be less encoded in individual neurons than the movement variables, except for the licking behavior (Engelhard et al., 2019; Musall et al., 2019; Stringer et al., 2019). We constructed a model with task variables, licking variables, and movement variables (upper body, nose, whisker, and mouth movements; these predictors were extracted from the video data recorded during behavioral tasks), and compared it with the model with task-related variables and licking variables (Figures S6A and S6B). The prediction accuracy of the latter model was not worse than the fitting of the full model in almost all imaged fields (Figures S6C and S6D). Thus, although we cannot exclude the possibility that some parts of the neuronal activity might reflect other movement variables more strongly than the cue and outcome variables, we considered that the neuronal activity strongly encoded the reward-predicting cue, current outcome, and recent outcome history.

We also applied multiple linear regression to the data obtained with wide-field calcium imaging (Figures S7A and S7B). In all areas, the prediction accuracy of the model steadily increased over the sessions (Figure S7C). From the early to late stage of learning, the weight of the current cue variable tended to increase in all areas, while that of the one-trial back outcome variable substantially increased in dmFrC, dlFrC, and M1 (Figure S7D). The weight of the current outcome variable was high in many areas in both stages of learning, while the weights of the one and two-trial-back cue variables and two-trial-back outcome variables remained small. These results suggest that the three dorsal frontal areas developed to strongly represent the reward-predicting cue and recent reward history through learning.

### Individual frontal cortical neurons do not simultaneously represent the reward-predicting cue with the current or past reward

Although the multiple linear regression model revealed the weight of each variable at each time-frame, it did not reveal the temporal pattern of each variable encoded by individual neurons, or whether or how multiple variables were represented by the same neurons. Therefore, we conducted a non-hierarchical clustering analysis to examine the combination of cue and reward variables that governed the activity of individual neurons in the four imaging areas. Neurons without any predictor variable with a P value of < 0.01 at any time-frame, except for lick-rate variables, were excluded from this analysis (2904 neurons were excluded). For the remaining neurons (14113 neurons), we used the Louvain-Jaccard clustering method to analyze the data for the neuron number × 606 dimensions (six cue and reward variables × 101 time-frames in each trial period). The Louvain-Jaccard clustering is a graph-based clustering method by community detection with modularity optimization (Blondel et al., 2008) and has been used to effectively process high-dimensional data such as the links of web pages, social network services (Chamberlain et al., 2018), and large-scale multicellular gene expression data (Macosko et al., 2015; Shekhar et al., 2016; Wu et al., 2017), without the premise of division number (see Methods for details). With this method, the remaining neurons were finally classified into 17 clusters.

Each cluster could be separately mapped and visualized in two-dimensional space by t-Distributed Stochastic Neighbor Embedding (t-SNE; Figure 8A). Each cluster showed a unique time-series of the weights of the cue and reward variables (Figure 8B). Clusters with high weights for the present cue (clusters 1 and 3) were well separated from the other clusters (Figure 8A). Any cluster did not have the combination of cue and outcome variables with high weights. By contrast, clusters 7 and 11 showed the combination of two outcome variables: the current outcome and recent reward history (Figure 8B).

**Figure 8.**
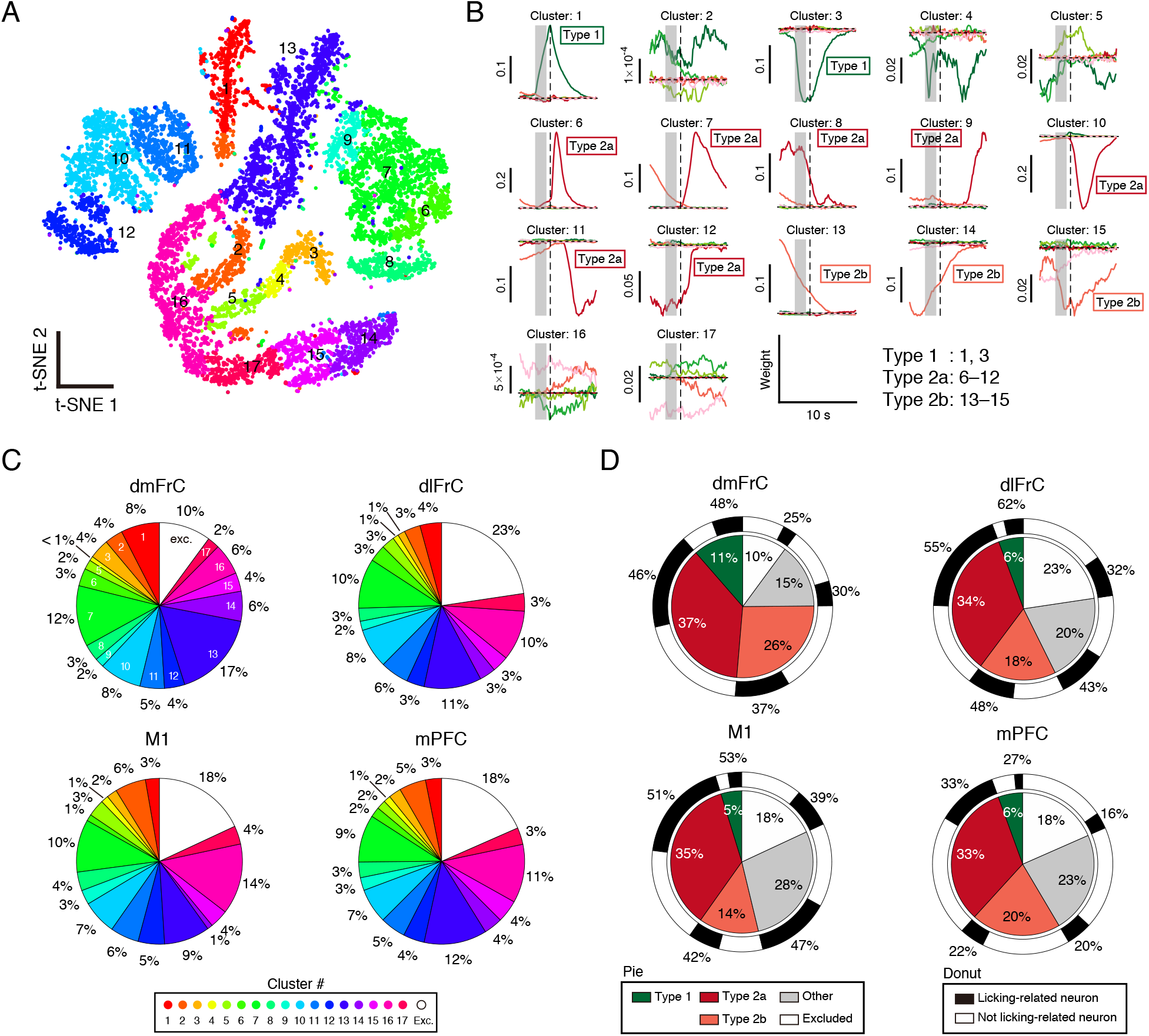
Unsupervised clustering of neurons according to the encoding modality. (**A**) Two-dimensional t-SNE map reflecting the coefficients of the task-related model (*α*_0_–*α*_2_ and *β*_0_-*β*_2_). Each dot corresponds to a single neuron. The colors indicate the cluster to which each neuron was assigned. Clustering analysis was performed independent of the t-SNE, and 17 clusters were finally obtained. Neurons without any task-related variable with P-values of < 0.01 at any time-frame were excluded from the clustering analysis (defined as “excluded”) and this t-SNE map. (**B**) Dynamics of the coefficients in each cluster. Green and red colors correspond to the mean weights of cue variables and reward variables, respectively, and deep, middle, and pale colors are those in the zero-, one-, and two-trials-back trials, respectively. SEM is not shown to facilitate clear data visualization. (**C**) Cluster ratios for each area. The number and color of each sector in the pie chart correspond with the clusters shown in (A). The white sector shows the proportion of excluded neurons. (**D**) Pie chart of type 1 (green), type 2a (red), type 2b (pink), other types (gray), and the excluded neurons (white). Black parts of the circumference indicate the lick-related neurons shown in Figure 5E. Clusters corresponding to each type are shown in (B).

For the sake of simplicity, the clusters were redefined according to the qualitative feature of the weight of each predictor variable as three specific types and ‘other types’ (Figure 8C and Table S1): type 1, a group that strongly represented the present reward-predicting cue (clusters 1 and 3); type 2a, a group that strongly represented the current outcome (clusters 6–12); type 2b, a group that strongly represented the recent reward history but not the current outcome (cluster 13–15); and other types, a group with small weights for predictor variables (less than the 95th percentile; clusters 2, 4, 5, 16, and 17). In mPFC, dlFrC, and M1, the sum of type 1, 2a, and 2b populations constituted 50–60% of the total, while in dmFrC neurons they formed 75% of the total. The proportions of types 1 and 2b were high in dmFrC compared with the other areas. The proportion of type 2a was similar (33–37%) among the four areas, and out of the type 2a predictor variables, the proportions of neurons with negative weights for the current outcome variable (clusters 8, 10, and 11) that were active in the reward omission were also similar (43–50%). The type 1, 2a, and 2b proportions were high at the depth of 500–800 μm (Figure S8). These results indicate that dmFrC neurons represented the reward-predicting cue and recent reward history more strongly than dlFrC, mPFC, and M1 neurons, while the neurons in all these areas similarly represented the current outcome.

Finally, we examined the proportion of licking-related neurons within each type of the neurons (Figure 8D). Licking-related neurons occupied 37% to 62% of each type of cluster in dmFrC, dlFrC, and M1, and 20% to 33% in mPFC. These proportions were similar to the proportions for all imaged neurons (Figure 5E). The Louvain-Jaccard clustering revealed that a majority of the frontal cortical neurons possessed task-related information, that the type of information each neuron possessed did not depend on whether the neuron was licking-related, and that the cue- or outcome-related information could be multiplexed with the licking-related information at the level of single frontal cortical neurons.

## Discussion

In the current study, we clarified changes in cortex-wide activity associated with the learning of a classical conditioning task, a simplified task that required stimulus-outcome association. The area-averaged activity remained highly correlated with the licking behavior in all dorsal cortical areas. This high correlation between movements and neuronal activities is consistent with recent reports using more complex behavioral tasks (Musall et al., 2019; Pinto et al., 2019; Stringer et al., 2019). Ongoing movements directly affect the interactions between the environment and animal; therefore, they should also affect the sensory and motor processing. Many dorsal cortical areas are related to this processing, and information regarding movements may therefore be broadly represented over these areas (Clancy et al., 2019; Goard et al., 2016; Pakan et al., 2018; Pinto et al., 2019). By contrast, mPFC had a smaller number of licking-related neurons than dmFrC, dlFrC, and M1. mPFC may process internal information (such as hippocampus-dependent memory, strategy switching, and conflict monitoring) more strongly than movement information (Euston et al., 2012; Friedman et al., 2015; Tervo et al., 2014).

The regression model suggests that a part of the cortex-wide activity evolved to possess information on the reward-predicting cue and recent reward history. The emergence of these information types was also detected in the striatum (Bloem et al., 2017). Irrespective of whether an appropriate choice is demanded or not, many cortical and subcortical areas would be able to use these types of information to adjust sensory attention and/or movement variability (Anderson, 2019; Pekny et al., 2015).

In the late stage of learning, dmFrC and dlFrC areas showed reward-predicting cue information after the cue presentation. dmFrC contained many neurons that positively and negatively represented the reward-predicting cue. The neurons with negative weights for the cue variable might be related to response inhibition (Aron et al., 2014), and may inhibit the anticipatory licking in cue B trials. The inactivation of these neurons by the photoinhibition of dmFrC might have disinhibited the anticipatory lick-rate in cue B trials, and as a result, the anticipatory lick-rate became similar between cue A and cue B trials (Figure 4E). In a dynamic foraging task, cue and action values were more strongly represented in dmFrC than in lateral sites (Sul et al., 2011). In the primate, the dorsomedial frontal cortex is assumed to transform stimulus values to motor commands (Cisek and Kalaska, 2005; Hare et al., 2011). Thus, dmFrC is probably critical for initiation of an appropriate motor action based on the value of a sensory stimulus (Siniscalchi et al., 2016, 2019).

The “dlFrC” used in our current study was originally defined as the ALM, in which intracortical micro-stimulation is known to induce tongue movement (Hira et al., 2015; Komiyama et al., 2010; Tennant et al., 2011). In fact, the dlFrC inactivation in the pretraining sessions is known to inhibit licking behavior, as recently demonstrated in VGAT-ChR2-EYFP mice (Bollu et al., 2019). Consistent with this, optogenetic inactivation of M1 did not inhibit licking in a Go-NoGo task (Allen et al., 2017). Although the representation of the reward-predicting cue during the cue period was weak in dlFrC, it was strong during the delay period (Figures 7A–7C). The neurons in the deep layer at 500–800 μm in both dmFrC and dlFrC had more information than those in the superficial layer (Figure 7A and S8C). These results are consistent with studies showing that layer 5 neurons in the ALM play more crucial roles in the planning and execution of movement than superficial-layer neurons (Chen et al., 2017b; Guo et al., 2017; Li et al., 2015, 2016). Considering these results together, we propose that the deep-layer dmFrC may be a critical site for accumulating reward-predicting sensory information and past and current reward information for the selection of appropriate actions, while the deep-layer dlFrC may be strongly related to retention of the reward-predicting sensory information that is required to execute certain types of selected movements.

The clustering analysis revealed three characteristic patterns of neuronal representation in individual neurons in the frontal cortex. First, licking information was simultaneously represented with the reward-predicting cue or the past and/or current reward information in 22–62% of the neurons in the four frontal cortical areas. This mixed representation of behavior-related and task-related information was also detected in the visual cortex, amygdala, and dopamine neurons in the ventral tegmental area (Engelhard et al., 2019; Gründemann et al., 2019; Stringer et al., 2019). It may be a general principle that ongoing movements affect the neuronal activity in a proportion of individual neurons during sensory processing. Second, a subset of neurons simultaneously represented the one-trial-back and current rewards (clusters 7 and 11). These neurons appeared to maintain high or low activity from the reward delivery to the end of the next cue presentation. The proportion of these neurons was similar among the four areas (14–17%). By contrast, the proportion of type 2b neurons without the current reward signal was highest in dmFrC.

Information regarding the recent reward history might be maintained by persistent activity by corticocortical connections and/or through the thalamus (Collins and Anastasiades, 2019; Guo et al., 2017; Rikhye et al., 2018; Tanaka et al., 2018). Third, in all clusters, the reward-predicting cue was not presented strongly with the past or current reward within individual neurons.

The highest proportion of type 1 neurons was detected in dmFrC. The type-1 neurons may send reward-predicting cue information to other dmFrC neurons that prepare the outcome-associated action in the decision-making process. Other areas including the orbitofrontal cortex, deeper mPFC than that imaged in the current study, and/or the striatum stably represent cue and action values, allowing update of the reward-predicting cue (Amarante et al., 2017; Aron et al., 2014; Bari et al., 2019; Ridderinkhof et al., 2004; Stalnaker et al., 2014; Zalocusky et al., 2016). These areas may directly or indirectly activate and/or inactivate type-1 neurons in dmFrC. Our results suggest that dmFrC acts as an initiator of cortex-wide neural ensembles related to reward prediction, and that the separate neuronal representations of task-related information in dmFrC, which were observed even in the condition without appropriate action selection, are the foundation for decision-making processes over time.

## Materials and Methods

### Animals

All animal experiments were approved by the Institutional Animal Care and Use Committee of the University of Tokyo, Japan. All mice were provided with food and water ad libitum, and housed in a 12:12 hour light-dark cycle (light cycle; 8 AM–8 PM). Male and female C57BL/6-derived transgenic mice (aged 2–8 months) were used for the wide-field calcium imaging experiments and photoinhibition experiments. The transgenic mice for wide-field imaging were obtained from homozygous Emx1-IRES-Cre mice (B6.129S2-Emx1^tm1(cre)Krj^/J, JAX stock #005628, Jackson Laboratory; Gorski et al., 2002) and hemizygous CaMKII-tTA::TITL-R-CaMP1.07 mice (directly obtained from Dr. F. Helmchen, they can now be purchased from Jackson laboratory as Ai143D, but they are not crossed with the CaMKII-tTA mice; Bethge et al., 2017). Transgenic mice for the photoinhibition were obtained from hemizygous VGAT-ChR2-EYFP mice (B6.Cg-Tg(Slc32a1-COP4*H134R/EYFP)8Gfng/J, JAX stock #014548, Jackson Laboratory; Zhao et al., 2011). Wild-type C57BL/6 mice (male, aged 2–3 months, SLC, Japan) were used for two-photon imaging.

### Surgical procedures

Mice were anesthetized by intraperitoneal or intramuscular injection of a ketamine and xylazine mixture (74 mg/kg and 10 mg/kg, respectively). After anesthesia, atropine sulfate (0.5 mg/kg) was administered intraperitoneally to improve breathing and bronchial secretion, and an eye ointment (Tarivid; 0.3% w/v ofloxacin, Santen Pharmaceutical, Japan) was used to prevent eye-drying and infection. During surgery, body temperature was maintained at 36–37°C with a heating pad. The head of the mouse was sterilized with 70% ethanol and the hair was shaved, and the skin covering the neocortex was incised. After the skull was exposed, the intermediate tissue on the skull was clearly removed. For wide-field imaging preparation, the temporal muscle was carefully dissected to improve the observation of temporal cortical areas. A custom head-plate (Tsukasa-Giken, Japan) was attached to the skull using dental adhesive (Estecem II, Tokuyama Dental, Japan). To prevent excitation light from directly entering through gaps between the head-plate and eyes of the mouse, the gaps were filled with a mixture of dental resin cement (Fuji lute BC; GC, Japan) and black-colored carbon powder (Wako chemical, Japan). To prevent drying of the skull surface, a thin layer of cyanoacrylate adhesive (Vetbond; 3M, MN) and dental adhesive (Super bond; Sun Medical, Japan) were applied. An isotonic saline solution with 5 w/v% glucose and the anti-inflammatory analgesic carprofen (5 mg/kg, Remadile; Zoetis, NJ) was injected intraperitoneally after all surgical procedures.

For the transcranial wide-field calcium imaging and photoinhibition experiments, the transparency of the layer of resin was generally low after the dental resin had cured, which affected the signal-to-noise ratio of the transcranial fluorescent imaging and the efficiency of photostimulation. Therefore, the surface of the resin was coated with a thin layer of nail polish, which was allowed to dry. This procedure was repeated a few times. In this manner, chronically stable optical access through the resin and skull was achieved (vessels in the brain surface were observed), and transcranial wide-field calcium imaging and photoinhibition could be performed.

For the preparation of two-photon imaging, viral injection was performed after fixation of the head-plate. Viral injections were performed with a pulled glass capillary (broken and beveled to an outer diameter of 25–30 μm; Sutter Instruments, CA) and a 5 μl Hamilton syringe (Cat. # 87930, Hamilton, NV). The glass capillary was filled with mineral oil (Nacalai Tesque, Japan) and the viral solution was drawn up from the tip just before the injection. The skull over the desired injection site was thinned with a 0.4 mm bar and a high-speed drill, and the glass capillary was inserted through the thinned bone. The stereotaxic coordinates used for the cortical injections were as follows: dorsomedial frontal cortex, dmFrC, AP +2.8 mm, ML −0.6 mm, DV 300–500 μm; dorsolateral frontal cortex, dlFrC, AP +2.5 mm, ML −1.8 mm, DV 300–500 μm; medial prefrontal cortex, mPFC, AP +2.8 mm, ML −0.6 mm, DV 1200 μm; primary motor cortex, M1, AP +0.5 mm, ML −1.2 mm, DV 300–500 μm. The AAV solution (AAV1-hSyn-NES-jRGECO1a, titer ~1 × 10^13^ GC/ml, the viral solution provided by Addgene, viral prep # 100854-AAV1; http://n2t.net/addgene:100854, RRID:Addgene_100854; original viral plasmid pAAV.Syn.NES-jRGECO1a.WPRE.SV40 was a gift from Douglas Kim and the GENIE Project) was infused over 20 minutes with a syringe pump (KDS310; KD Scientific, MA). To prevent backflow and improve the spreading of solution, the glass capillary was left in place for over 10 min after completion of the injection. After ~30 min from the start of the injection, the glass capillary was carefully withdrawn. For the injection to mPFC, to minimize background fluorescence from solution backflow through the space made by the glass capillary insertion, the axis of the glass capillary was angled 30–40° from the horizontal plane. After the injection, the injection site was covered with silicone elastomer (Kwik-cast, World Precision Instruments, FL) and then dental acrylic.

Two to three weeks after injection, surgery was performed to implant a glass window for two-photon imaging. Mice were anesthetized by isoflurane (Pfizer, NY; 3–4% for induction and ~1% during surgery) and their body temperature was maintained at 36–37°C with a heating pad. To prevent cerebral edema, dexamethasone sodium phosphate (1.32 mg/kg; Decadron, Aspen Japan, Japan) was administered intraperitoneally 30 min before the start of surgery. As in Goldey et al., (2014), the glass window consisted of two square cover slips (for dmFrC/dlFrC/mPFC, No. 1, 0.12–0.17 mm thickness and 3 mm square; and No. 5, 0.45–0.60 mm thickness and 2 mm square; Matsunami Glass, Japan) or two circular coverslips (for dmFrC/dlFrC/mPFC, No. 1, 0.12–0.17 mm thickness and 2.5 mm diameter; and No. 5, 0.45–0.60 mm thickness and 1.5 mm diameter; for M1, No. 1, 0.12–0.17 mm thickness and 3.0 mm diameter; and No. 3, 0.25–0.35 mm thickness and 2.0 mm diameter; Matsunami Glass), which were glued together with UV-curing optical adhesive (NOR-61; Norland Products, NJ). The geometry of the glass coverslips did not clip excitation light exiting from the front of the objective (described in detail in Kondo et al., 2017). After the head was sterilized, the dental resin and silicone elastomers above the desired observation site were removed, and the skull surface was cleaned with artificial cerebrospinal fluid (aCSF). Craniotomy was performed with a 0.4 mm bar and high-speed drill. The size and geometry of the craniotomy was determined to match the implanted glass window. To minimize heat damage during drilling of bone, aCSF was administered to the operative zone. A glass window was placed over the craniotomy and the edge was sealed with cyanoacrylate adhesive, dental resin cement, and dental adhesive. The dura mater was left intact as far as possible; however, in some cases (e.g., for deep imaging preparation, where the dura was opaque and too thick) it was removed with great care and attention. After implantation of the glass window, 250 μl saline with the anti-inflammatory analgesic carprofen were administered intraperitoneally. The mice were then returned to their home cage and allowed to rest for at least 1 day before the imaging sessions commenced.

### Classical conditioning task

Before starting behavioral training, mice were placed under a water-deprived condition in their home cage. Food was provided ad libitum. During the experiments, their body weight was controlled at 80–85% of the initial weight. If body weight had not sufficiently increased after behavioral training, an appropriate volume of water or agar gel was given. On the rest day (typically weekends), an agar gel corresponding to the daily water consumption was also given. During the conditioning sessions, mice were head-fixed via their head-plate and placed in a body chamber. The sessions were carried out once per day, with a duration of 40–60 min.

The mice were submitted to a classical conditioning paradigm, except that the reward (unconditioned stimulus, US) was delivered in a probabilistic manner, depending on a presented cue-tone (conditioned stimulus, CS). A waterspout was placed in front of their mouth, and a drop of water (4 μl) was given as the US (reward). The mice were able to freely lick at any time, with licking being monitored by an infrared LED sensor or a touching detection circuit. One of two sound cues (6k or 10k Hz) was presented for 1.5 s as the CS, and after a delay of 0.5 s, the US was delivered at a probability corresponding to the sound cue. Inter-trial-intervals were 8 ± 2 s. The reward probability was set at 0.7 for one of the sound cues and 0.3 for the other. Either sound was randomly presented in each trial. In 2 or 3 days of pretraining prior to starting the reward probability with conditioning, the mice experienced a general classical conditioning, in which the reward was always delivered after either cue. For the conditioning with different probabilities of reward delivery, when the lick-rate during the sound cue onset to the reward delivery (2 s) for the high reward probability was greater (P-values of < 0.01, one-tailed Wilcoxon rank-sum test) than that for the low reward probability in three consecutive sessions, we considered the training to be completed. Wide-field calcium imaging was performed daily from the start to end of training. Two-photon calcium imaging and photoinhibition experiments were conducted after the training was completed.

### Photoinhibition of cortical areas

Blue light for activating channelrhodopsin-2 (ChR2) in inhibitory neurons was supplied from a laser diode module with an SMA optical fiber connector (BioRay FR, 450 nm, 50 mW, Coherent, CA). The laser module was operated by a StingRay laser controller (Coherent, CA), and the output intensity and timing were controlled by the analog-voltage output from a NIDAQ system. The laser beam was delivered through an AR-coated multimode optical fiber (M50L02S-A, Ø50 μm, 0.22 NA, 400–700 nm, Thorlabs Inc., NJ) and a fiber port (PAF2S-7A, f = 7.5 mm, 350–700 nm, Ø1.23 mm Waist, Thorlabs Inc., NJ) to a two-dimensional galvanometric scanning system (part of a laser scanning microscope, Olympus, Japan). The fiber port and scanning system were connected via a CNC-machined laboratory-made adaptor. The laser beam was focused on the brain surface (350 μm diameter at 4σ of Gaussian profile) through an achromatic doublet lens (AC254–200-A-ML, f = 200 mm, 400–700 nm, Thorlabs Inc., NJ).

Photoinhibition was performed on dmFrC (AP +2.8 mm, ML 0.5 mm from bregma), dlFrC (AP +2.5 mm, ML 2.0 mm from bregma), M1 (AP +0.5 mm, ML 1.0 mm from bregma), and the primary somatosensory barrel area (S1b, AP −1.2 mm, ML 3.2 mm from bregma). The photoinhibition trials were determined with a probability of 50%, being independently set according to the type of CS, and the order of the inhibited locations was randomized. The illumination order was reset every time the illumination was completed in four locations. The photoinhibition was performed bilaterally by alternately going back and forth between the same regions in both hemispheres. The photoinhibition train was constructed using rectangular pulses (40 Hz/hemisphere, duty-cycle 0.9, time-averaged intensity 6–9 mW/hemisphere) with a pulse train duration of 3 s and illumination from the cue onset. To suppress rebound activities after photoinhibition (Guo et al., 2014), the laser intensity was linearly decreased over the last 500 ms of the stimulus train. In the control and photoinhibited conditions, to prevent behavioral change from occurring when the photoinhibition light entered the eyes of the animal, a ‘masking flash’ was provided in all trials using a blue LED array placed in front of the animal (10 Hz, same duration and onset as the laser illumination). Animals were habituated to the masking flash prior to the photoinhibition, and it was maintained until the classical conditioning learning was completed.

To estimate the effect of photoinhibition on cue discriminability and licking, the discrimination score (DS) and modulation score (MS) were defined using the area under the receiver operating characteristics curve (auROC). The definitions of DS and MS are as follows:

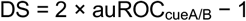

where auROC_cueA/B_ was calculated from the lick-counts during the cue presentation and delay periods when cue A and B were presented.

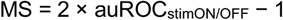

where auROC_stimON/OFF_ was calculated from the lick-counts during the cue presentation and delay periods when the photoinhibition was performed. Therefore, if DS is close to +1 or −1, it indicates that the mice are good ‘discriminators’ of cue A or B according to their licking. Conversely, if DS is close to 0, it indicates that the mice are bad ‘discriminators’ of the two cues. Similarly, the closer MS is to +1 or −1, the more the lick-count is respectively increased or decreased by the photoinhibition, and values close to 0 indicate that the lick-count was little-changed by the photoinhibition.

### Wide-field calcium imaging

Wide-field imaging was performed with Emx1-Cre::CaMKII-tTA::TITL-R-CaMP1.07 mice subjected to the transcranial preparation. Image acquisition was performed with a fluorescent zoom macroscope equipped with a high-numerical aperture macro lens (Plan-NEOFLUAR Z 2.3×, NA 0.57; Carl-Zeiss, Germany) and an EM-CCD camera (iXon Ultra 888; Andor Technology, UK). Images were captured by a Solis camera system (Andor Technology, UK) using a sampling-rate of 30 frames/s, pixel resolution of 512 × 512 pixels, and 2 × 2 binning. Optical zoom magnification was set to cover the entire dorsal cortex in each mouse. An excitation light source (HXP200C; Carl-Zeiss, Germany) with a filter-set (IX3-FGWXL, excitation, 530–550 nm; dichroic, 570 nm; emission, 575 nm LP; Olympus) and barrier filter (FF02-617/73-25, 580–655 nm; Semrock, NY) was used to observe R-CaMP1.07 fluorescence. A total of 54000 imaging frames were acquired per session (30 min/session). No apparent fading of the fluorescent signal was observed in the experimental condition across days.

### Two-photon calcium imaging

Two-photon imaging was conducted using an FVMPE-RS system (Olympus, Japan) equipped with a 25× water immersion objective (for imaging of the mPFC and dmFrC: XLPLN25XSVMP2, Olympus; for imaging of the dlFrC and M1: XLPLN25XWMP2, Olympus) and a 1100 nm excitation light source (Insight DS +Dual, Spectra Physics, CA). Fluorescence emissions were collected using a GaAsP photomultiplier tube (Hamamatsu Photonics, Japan). To improve optical access to the deep tissue, the back aperture of the objective was underfilled with a diameter-shortened (7.2 mm, in comparison with the back aperture of 14.4 mm or 15.1 mm) laser beam. In this procedure, the effective excitation NA was calculated to be roughly 0.5. The details of the optical assessment were described previously (Kondo et al., 2017). To make the angle of incidence between the excitation light and the plane of the glass window vertical, the angle of the task device and head-fixing apparatus was finely tuned using biaxial goniometers. The imaging parameters included a frame acquisition rate of 30 frames/s, a pixel dwell time of 0.067 μs, an imaging field size of 512 × 512 pixels (0.994 μm/pixel) or 512 × 160 pixels, and three-frame averaging to increase the signal-to-noise ratio in some of the experiments to image the mPFC. The duration of one imaging session was 15 min, and two imaging sessions from different depths were performed in each daily experiment, unless otherwise noted. For each mouse, imaging was conducted for 2–5 days.

### Behavioral video monitoring

Two high-speed video cameras (Scout scA640-70gm, Basler, Germany, and a fixed-focus lens M0814-MP, focal length, 8 mm, Computer, NC) were used to synchronously capture the upper parts of the bodies and faces of the mice. Using a synchronization signal from the microscopy, the image acquisition rate of the high-speed cameras was synchronized to that of the microscopy. An infrared LED light source (940 nm, 60 LEDs) was positioned at the back of each camera.

### Image processing for wide-field calcium imaging data

We analyzed the data from six mice that showed P-values of < 0.05 for the difference in anticipatory lick-rate between cue A and cue B trials. All analyses were conducted using custom Matlab routines (R2016a, R2018a, MathWorks, MA). Each image stack was rigidly aligned to the top view of the Allen Common Coordinate Framework version 3 (Allen CCF) using 10 control points. The control points were set at the bilateral anterior nexus of the olfactory bulbs, center and side tips of the frontal blood vessels, lambdoid suture, bilateral posterior tips of the dorsal cortex, and bilateral side tips of the dorsal cortex. After image registration, a binary mask was made from the Allen CCF (a contour of the top view), and values outside of the mask were set to zero to exclude fluorescent changes in the pixels outside of the cerebral cortex. To reduce computational costs and time, an image clipping procedure was applied to the image stack, and the pixels of the left hemisphere were used for the following analyses. The singular value decomposition (SVD) of the clipped image stack was calculated to reduce dimensionality and noise. From the result of the SVD, the spatial component *U* (of size pixels × components), singular value S (of size spatial components × temporal components), and transposed temporal components *V*^T^ (of size components × frames) were obtained. The top 12 singular values were used to reconstruct the original dimensions of the image stack, with these 12 singular values and their spatiotemporal components explaining 99.93% of the variance of the original. Using the reconstructed data, the fluorescent intensity change (ΔF/F) was calculated for each pixel with the 10th percentile value in a ±15 s-interval around each frame. For analyses using ROIs, twelve ROIs (10 × 10 pixels, rectangular shape) were set for the brain areas described below, with the stereotaxic coordinates being defined according to the top view of the Allen CCF and previously reported areas, with these being anatomically and functionally identified: dmFrC (AP +2.8 mm, ML 0.8 mm); dlFrC (AP +2.5 mm, ML 2.0 mm); M1 (AP +1.2 mm, ML 1.0 mm); primary somatosensory forelimb area (S1FL, AP +0.4 mm, ML 2.2 mm); primary somatosensory cortex hindlimb area (S1HL, AP −0.5 mm, ML 1.5 mm); primary somatosensory cortex mouth area (S1m, AP +1.4 mm, ML 3.0 mm); primary somatosensory cortex nose area (S1n, AP 0 mm, ML 3.4 mm); primary somatosensory cortex barrel area (S1b, AP −1.2 mm, ML 3.0 mm); primary auditory cortex (Aud, AP −2.5 mm, ML 4.0 mm); primary visual cortex (Vis, AP −4.0 mm, ML 2.2 mm); posterior parietal cortex (PPC, AP −2.0 mm, ML 2.0 mm); retrosplenial cortex (RSC, AP −2.4 mm, ML 0.5 mm). The stereotaxic coordinates indicate the center positions of the ROIs. For each session and mouse, the ROI positions were manually fine-tuned. The fluorescence intensity of each ROI was determined by averaging the included pixels.

### Image processing for two-photon calcium imaging data

Raw image sequences acquired with the FVMPE-RS system were loaded into MATLAB using custom-written scripts (Oir2stdData, https://github.com/YR-T/oir2stdData). Motion correction for image stacks was performed using the NoRMCorre algorithm (Pnevmatikakis and Giovannucci, 2017, https://github.com/flatironinstitute/NoRMCorre). After performing rigid motion correction, the aligned image stack was further corrected with a non-rigid motion correction method. After this motion correction, the images were three-frame averaged before being analyzed. A constrained non-negative matrix factorization (cNMF) algorithm was employed to extract neuronal activities from a time-series of images (CaImAn package for MATLAB, Pnevmatikakis et al., 2016, https://github.com/flatironinstitute/CaImAn-MATLAB). Automatically extracted neuronal activities that did not reach the following criteria were removed: maximum size of spatial components, 250; minimum size of spatial components, 30; threshold of the spatial correlation between the estimated component and raw data, 0.6; threshold of the temporal correlation between the estimated component and raw data, 0.5; minimum signal-to-noise ratio of the estimated component, 0.2. Finally, using a pre-trained convolutional neural network discriminator (version Feb. 13, 2018), those ROIs with a pseudo neuropil structure (dendrites and/or axons) were removed (network responses with a confidence level of > 0.2 were accepted). The detrended relative fluorescence changes (ΔF/F) were calculated for eight percentile values over an interval of ± 30 s around each sample time point (Dombeck et al., 2007).

### Behavioral video analysis

Videos obtained from the high-speed camera were processed with FaceMap package (Stringer et al., 2019; https://github.com/MouseLand/FaceMap) to identify the mouth, nose, whiskers, and upper body parts. ROIs were set on the forelimbs and thorax, nose, mouth, and whisker-pad, and the SVD of the motion-energy (absolute frame-to-frame difference) of each ROI was calculated. The software generated “mot¡on-mask”, which was calculated from the spatial component *U* and the singular value S found by SVD, and the time-series of separable movements in singular value space were calculated by multiplying the motion-mask by the original time-series data. Most of the dominant movements of motion-energies in each ROI (body parts) were used in the regression analysis.

### Neuronal activity analysis for wide-field imaging data

Neuronal activities extracted from the ROIs were converted into Z-scores and aligned to the onset of each cue-tone. To estimate the activities during learning stages, the neuronal activities in each session were averaged for each ROI and for each leaning stage (early; session 1–2, middle; session 4–5, late; session 7–8). To examine how the neuronal activities and licking behavior changed during the 2 s period from the cue-tone onset to the reward delivery timing, the neuronal activities during this period were divided into 200 ms bins. The neuronal activity of each time-bin was compared with the baseline activity for the 0.5 s before the cue-tone onset. We performed factor analysis to visualize the cortex-wide neuronal trajectory (Harvey et al., 2012; Makino et al., 2017). Neuronal activities were decomposed into five factors with the *factoran* built-in Matlab function. The distance of the neuronal trajectory between cue A and cue B trials was defined by Euclidean distance in a variable space obtained from the factor analysis in each learning stage. When the relationship between the neuronal trajectories and licking was examined, pairwise trial-by-trial correlations of lick-rate were binned for every 0.2 from −1 to 1.

It was reported that in mice expressing a genetically-encoded calcium indicator knocked-in to the TIGRE-locus and driven by Cre-recombinase under the control of pan-neuronal promoter (e.g., Emx1 promoter), hyperexcitability was invoked in the frontal cortex and propagated to the entire dorsal cortex (Steinmetz et al., 2017). To check whether this phenomenon was observed in Emx1-Cre::CaMKII-tTA::TITL-R-CaMP1.07 transgenic mice, spontaneous activities were evaluated in five typical ROIs (MOs, MOp, S1b, RSP, Vis). For periods of 20–30 min, spontaneous activities under awake and resting-state conditions were measured in four mice. After the data were pre-processed using the same procedure as described above, square ROIs were set. A peak detection (*findpeaks*, a built-in function of the signal processing toolbox in Matlab) was performed for the time-series of neuronal activities obtained from each ROI. Aberrant activities with large amplitudes and short-durations (Steinmetz et al., 2017) were not observed in our preparation (Figure S1).

We performed Granger causality analysis using the multivariate Granger causality toolbox (Barnett and Seth, 2014) and adopted the procedures previously described in Makino et al., 2017. We used the pairwise conditional Granger causality as the ‘causality’. The conditional causality was defined by the elements of ‘to’ (X) and ‘from’ ROIs (Y), and for a given X, conditional causality for a given Y is the degree to which the past activity of Y helps to predict the time-series of X activity, over the degree to which X is predicted by the past activities of X and the other conditional ROIs. For each ROI, *ΔF/F* during the cue presentation and delay periods was subtracted by baseline activity (the mean activity for 1 s before the cue onset) to satisfy the covariance stationarity. Trial-by-trial neuronal activity in each ROI was then concatenated. Time-domain Granger causality was then computed from the concatenated time-series with the multivariate Granger causality toolbox. This toolbox computes Granger causality based on vector autoregressive modeling. The model order (the number of time-lags) was estimated by Akaike’s Information Criterion with the maximum set to be 20. Ordinary-least-squares was used to compute regression coefficients. Pairwise conditional causality was then calculated and the causality which had statistical significance (P < 0.01, false discovery rate [FDR] corrected) were only kept and others were set to zero. The median of the causality across mice was obtained at each learning stage.

### Neuronal activity analysis for two-photon imaging data

The neuronal activity that was automatically extracted with the cNMF algorithm was converted to Z-scores and aligned to the onset of the cue presentation. To find the neuronal activity associated with the licking behavior, cue, reward delivery, reward omission, and reward anticipation (Wagner et al., 2017), and to make statistical comparisons in the size of the activity between different periods, we used the following four criteria. First, neurons that showed a significant (P value < 0.05) trial-averaged correlation coefficient of > 0.3 between the lick-rate and activity trace were assigned as ‘licking-related neurons’. To calculate the correlation coefficient, both types of cue-tone presentation (A and B) and both types of reward delivery (acquisition and omission) were used. In the following classification, the licking-related neurons were removed. Second, neurons in which the average activity during 1.5–2.0 s was larger than that during 0–0.5 s by ≥ 0.3 standard deviation (s.d.), the rewarded-trial-average activity during 1.5–2.0 s was smaller than that during 2.0–2.5 s by ≤ 0.2 s.d., and the unrewarded-trial-average activity during 2.0–2.5 s was larger than that during 1.5–2.0 s by ≥ 0.2 s.d., were assigned as “reward anticipation neurons”. Third, among the neurons that did not satisfy the criteria 1 and 2, those in which the rewarded-trial-average activity during 2.0–3.0 s was larger than that during 1.0–2.0 s by ≥ 0.3 s.d., and the unrewarded-trial-average activity during 2.0–3.0 s was smaller than that during 1.0–2.0 s by ≤ 0.2 s.d., were assigned as “reward acquisition neurons”. Conversely, those in which the unrewarded-trial-average activity during 2.0–3.0 s was larger than that during 1.0–2.0 s by ≥ 0.3 s.d., and the rewarded-trial-average activity during 2.0–3.0 s was smaller than that during 1.0–2.0 s by ≤ 0.2 s.d., were as “reward omission neurons”. Fourth, among the neurons that did not satisfy criteria 1–3, those in which, in one of each of the cue presented trials, the average activity during 0–1.0 s was larger than that during −1.00 s by > 0.3 s.d., were assigned as “cue neurons”.

### Neuronal encoding model

We conducted multiple linear regression analysis to investigate how the task-related events (cue A or B, and reward delivery or no-delivery) and their histories were encoded in the neuronal activity. Our primary purpose in this analysis was to estimate the trial-averaged encoding in a single neuron from the trial-by-trial variabilities of task-related events and licking behavior. Therefore, we performed a frame-by-frame regression using predictors that had trial-by-trial variance, but not a linear regression with the convolution of many kernels and task-related events in the time-dimension (e.g., Engelhard et al., 2019; Musall et al., 2019; Pinto and Dan, 2015). To handle the licking behavior in this manner, the licking variables were binned into separable time windows for each trial.

The neuronal activity at the *i-th* time-frame was regressed according to the following formulation:

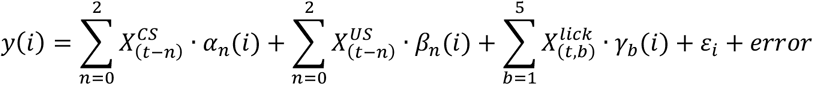

where *X*^CS^(*t*) is 1 or −1 if cue-tone A or B are presented in the t-th trial, respectively; *X*^US^(*t*) is 1 or −1 if the reward is delivered in or not in the t-th trial, respectively; *X*^lick^(*t,b*) is the mean of the z-scored lick-rate in each 2 s bin of the t-th trial (*b* = 1, −2–0 s from the onset of the cue presentation, *b* = 2, 0–2 s; *b* = 3, 2–4 s; *b* = 4, 4–6 s; *b* = 5, 6–8 s); *α*_*n*_(*i*), *β_n_*(*i*), and *γ_b_*(*i*) are the weights of *X*^CS^_(*t-n*)_, *X*^US^(*t-n*) and *X*^lick^(*t-n*), respectively; and *ε_i_* is a bias term. We also included the cue and reward information in one (*n* = 1) and two (*n* = 2) trials before the current trial evaluated using the formulation. The number of all time-frames was 101 (from before 2 s to the cue onset to after 8 s, 10 frames/s). The effect-size was set to the weight value of the predictor variable if its P value was less than 0.01, and otherwise it was set to zero.

The model performance was estimated by 5–fold cross-validation in each time-frame. The linear regression was performed by randomly allocating 80% of the trials to a training set, and the model response was then estimated using the residual 20% of trials (the test set). The metric for estimating the model fitness was the squared Pearson’s correlation R between the actual neuronal activity and the concatenated model responses, referred to as the cross-validated *R*^2^ (cv*R*^2^, Figure S4B). The five cv*R*^2^s values obtained from the 5-fold cross-validation were averaged, and the averaged values were used for further analyses.

When 5 surrounding time-frames (that is, 100 ms-bin) of the lick-rate were used for the regression at each time-frame, cvR^2^ was lower than that when the five 2 s-bin licking variables was used described above in all cortical areas and depths (100–300 μm of dmFrC, 0.173 ± 0.101 vs. 0.226 ± 0.115; 300–500 μm of dmFrC, 0.187 ± 0.0928 vs. 0.267 ± 0.104; 500–800 μm of dmFrC, 0.225 ± 0.112 vs. 0.287 ± 0.121; 100–300 μm of dlFrC, 0.177 ± 0.0881 vs. 0.252 ± 0.103; 300–500 μm of dlFrC, 0.167 ± 0.0783 vs. 0.254 ± 0.0909; 500–800 μm of dlFrC, 0.217 ± 0.133 vs. 0.272 ± 0.130; 100–300 μm of M1, 0.139 ± 0.0934 vs. 0.177 ± 0.0957; 300–500 μm of M1, 0.133 ± 0.0719 vs. 0.186 ± 0.0832; 500–800 μm of M1, 0.136 ± 0.0637 vs. 0.196 ± 0.0783; mPFC, 0.149 ± 0.0911 vs. 0.210 ± 0.0932; in all areas and depths, the P-values were almost zero [< 2.23 × 10^-308^] with Wilcoxon sign-rank test, FDR corrected). Thus, we used the five 2 s-bin licking variables as the predictor variables for licking.

A multiple linear regression analysis was similarly performed for the wide-field calcium imaging data. In each session for each mouse, the neuronal activity in each of the 12 ROIs was regressed with three cue variables, three reward variables, five licking variables, and a bias term. The performance of the model was also estimated in a similar way to that of the two-photon imaging data.

To examine whether the task-related variables (three cue and outcome variables) may be less encoded in individual neurons than the movement variables, except for the licking behavior, we compared the task-encoding model with the task and lick variable-sets, and the full model with the task, lick, and video variable-sets, where a licking variable-set consisted of 2 s averaged lick-rates in the current trial (five variables) and a video variable-set consisted of the current 2 s averaged trial, most dominant motion-energy calculated from four ROIs (five in a single ROI × 4 ROIs = 20 variables). To obtain the values of cvR^2^ in each model, 5-fold cross-validation was performed.

#### Clustering analysis

As each predictor variable had 101 time-frames, the 606-dimensional data (six task variables × 101 time-frames) was clustered with the following procedures, as may be applied to a large RNA-sequencing dataset (Chen et al., 2017a; Levine et al., 2015; Macosko et al., 2015; Shekhar et al., 2016).

#### Dimensionality reduction

We performed principal components analysis (PCA) on the data from all neurons, with 606 regression coefficients. Each neuron was classified into a cluster using the regression coefficients (weights) of the six predictor variables except for the five lick-rate variables. Neurons that did not have any time-frame and with one or more regression coefficients (P-values of < 0.01) for any of the cue and reward variables except for the lick-rate variables and bias term were excluded from the clustering analysis (564, 1411, 555, and 374 neurons from the pooled data of dmFrC, dlFrC, M1, and mPFC, respectively). Seven dimensions with a significant contribution (P < 0.01) in the PC space were used for further analysis. P-values were calculated by permutation test with circular shifting, with the permutation being repeated 100 times.

#### Visualization

The seven-PC-dimensional data were mapped to a two-dimensional space by t-distributed Stochastic Neighbor Embedding (t-SNE). We used an FFT-accelerated interpolation-based t-SNE package (FIt-SNE, Linderman et al., 2019; Maaten, 2014, https://github.com/KlugerLab/FIt-SNE) for the calculation. The hyper-parameters used were: numbers of iterations = 3000, perplexity = 30, and early exaggeration = 16. The mapping to the two-dimensional space was used only for visualization, not for clustering analysis.

#### Clustering

Louvain-Jaccard clustering of PC space data was performed. Louvain-Jaccard clustering involves a recently proposed algorithm for network analysis, such as analysis of social networks (Chamberlain et al., 2018), and performs clustering using the similarity between nodes in a graph structure. Recently, it was reported that Louvain-Jaccard clustering allowed large-scale data obtained by RNA-sequencing to be reliably clustered on the basis of the amount of expressed genes (Shekhar et al., 2016; Wu et al., 2017).

The calculations with this algorithm consisted of the following sub-steps: 1. Generation of a K-nearest neighbor (KNN) graph. Determination of a graph structure in which each node was connected to the K-nearest neighbor points (K = 30) in the PC-dimensional space. 2. Calculation of the Jaccard similarity index (JI) of each node in the KNN graph and determination of the weights according to the calculated JIs. 3. Performance of Louvain clustering on the KNN graph weighted by JI. The number of clusters was determined by maximizing the modularity Q. The metrics of the edges inside communities were compared with the edges between communities at a certain community-segmentation. For these procedures, we used the genLouvain MATLAB implementation of the Louvain clustering algorithm (Lucas G. S. Jeub, Marya Bazzi, Inderjit S. Jutla, 2011; Mucha et al., 2010), which resulted in 21 clusters.

#### Cluster merging

In order to merge over-split clusters, cluster merging was performed according to the method of Shekhar et al., 2016 with some modifications. The following process resulted in a final number of 17 clusters.

For each cluster (cluster N), the nearest and second nearest neighbors of the cluster (clusters M_1_ and M_2_, respectively) were obtained according to the distances defined as 1 – Pearson’s correlation coefficient in PC space. To determine the number of differentially expressed (DE) coefficients between cluster N and cluster M_1_ or M_2_, the following calculations were performed; First, for each of the 606 coefficients (six variables × 101 time-frames), we calculated the percentage of neurons in which the coefficient showed a significant contribution (P < 0.01) to actual neuronal activity. These percentages were defined as *P*_N_, *P*_M1_, and *P*_M2_ in clusters N, M_1_, and M_2_, respectively. In cluster N, *P*_N_ was *n_c_*/*n*, where *n_c_* was the total number of neurons with the significantly contributed coefficient in cluster N and *n* was the total number of neurons in the cluster N. If *P*_N_/*P*_M1_ (or *P*_M2_) was < 2, *P*_N_ was < 0.2, or *P*_M1_ (or *P*_M2_) was < 0.2, the coefficient was not used for the calculation of the differential expression between the clusters N and M_1_ (or M_2_). Next, for each coefficient, we determined whether *P*_M1_ (or *P*_M2_) was significantly larger than *P*_N_ by the binomial test as P_binomial_ = 1 – *binocdf*(*n_c_, n, P*_M1_ [or *P*_M2_]), where *binocdf* is a MATLAB built-in function in the statistics and machine-learning toolbox. The P_binomial_ of all the coefficients obtained using the methods described above were converted to FDR using the Benjamin and Hochberg method (Benjamini and Hochberg, 1995).

Although gene expression levels such as transcriptome data are value of ≥ 0, the coefficients in this study could take positive or negative values, so the difference when a coefficient was positive value in one cluster and negative value in another one could not be detected with the binomial test (as the binomial test considers the probability of whether certain phenomena occur or not; for example, whether a gene expresses in two cells). Therefore, we examined whether the distribution of values of the significantly contributed coefficients was similar between cluster N and M_1_ (or M_2_) by Wilcoxon rank-sum test (P_ranksum_ < 0.01, FDR corrected). If the P_ranksum_ was < 0.01, the corresponding FDR was used for determining the differentially expression. Otherwise, the coefficient was not used for the calculation of the differentially expression between clusters N and M_1_ (or M_2_).

Finally, the coefficients which had the FDR < 0.01 were defined as the DE coefficients between clusters N and M_1_ (or M_2_). The total number of the DE coefficients was less than 30, the cluster-pair was merged into one cluster. If both cluster-pairs (N and M_1_, and N and M_2_) had < 30 DE coefficients, the pair which have a closer distance was merged. The same process was repeated until there was no cluster M1 or M2 reaching the above criteria.

#### Cluster sorting and grouping by the qualitative feature

After performing the cluster merging, we sorted the 17 clusters with the “encoding metrics” calculated by following the procedure for the comprehensive clarity. The metrics contains “magnitude” and “sign” of the coefficients in each cluster (Table S1). 1) For each of the six predictor variables in each neuron, the absolute values of the coefficients at 101 time-frames were summed. In addition, to sort the clusters according to positive to negative encoding order for each variable, the “sign” was determined by trapezoidal numerical integration of the raw value of the 101 coefficients for each neuron. 2) The neuron-averaged “magnitude” and “sign” of each variable was calculated in each cluster. 3) In each cluster, the largest “magnitude” of the variable was determined. The sorting procedure described below was applied using the clusters with the largest variable in the order of *α*_0_ to *α*_2_, then *β*_0_ to *β*_2_. 4) For *α*_0_, by using the neuron-averaged “magnitude” and “sign” of each cluster, the clusters with a positive “sign” on average were sorted in descending order of “magnitude”. Next, the same sorting was performed for the clusters with a negative “sign”. 5) Step 4 was repeated for all other variables. 6) To classify whether the neurons in the cluster adequately encoded the task events in six predictor variables, the neuron-averaged “magnitudes” were averaged over the six variables as task-encoding magnitude in each cluster. If the task-encoding magnitude of the cluster was smaller than the 95th percentile of the 6-variable-averaged magnitudes from 14113 neurons (5.3379), the cluster was excluded from the feature-based re-grouping.

Finally, by using the sorting result and the encoding metrics of each cluster, a qualitative feature-based grouping of clusters was performed. Each cluster was assigned to one of four groups: type 1, a group that strongly represented the present cue; type 2a, a group that strongly represented the current outcome; type 2b, a group that strongly represented the recent reward history, but not the current outcome; the other type, a group with small weights of predictor variables (less than the 95th percentile). The encoding metrics used in this cluster sorting and the feature-based grouping are shown in Table S1.

### Statistics

The data are presented as mean ± standard deviation, and the error bars in graphs represent the standard error of the mean unless otherwise noted. The Wilcoxon rank-sum test, signed-rank test, paired t-test, Friedman test with post-hoc multiple comparison (correction with dunn-sidak method), Pearson’s correlation tests, binomial test, and permutation tests described above were used for statistical comparisons. Pairwise comparisons were two-tailed unless otherwise noted. To compensate for multiple comparisons, the false-discovery rate (FDR) was calculated using the Benjamin and Hochberg method (Benjamini and Hochberg, 1995). No statistical tests were run to predetermine the sample size, and blinding and randomization were not performed.

### Data and code availability

Oir2stdData, the Matlab program directly imports from Olympus.oir files, is available at https://github.com/YR-T/oir2stdData. All raw-data and the custom-written code used in this study are available from the lead contact Masanori Matsuzaki upon reasonable request.

## Supporting information

Suppl. figre and table and the legend

## Author Contributions

Conceptualization, M.K. and M.M.; Methodology, M.K.; Software, M.K.; Formal Analysis, M.K.; Investigation, M.K.; Data Curation, M.M.; Writing – Original Draft, M.K. and M.M.; Writing – Review & Editing, M.K. and M.M.; Visualization, M.K.; Supervision, M.M.; Project Administration, M.M.; Funding Acquisition, M.K. and M.M.

The authors declare no conflict of interest.

## Supplemental information

The Supplemental information includes 8 figures, 1 table, and 1 video.

## Acknowledgments

We thank Dr. F. Helmchen for providing CaMKII-tTA::TITL-R-CaMP1.07 mice; Y. Hirayama and M. Nishiyama for animal care and breeding; Dr. Y.R. Tanaka for statistical modeling; Dr. S.I. Terada for development of optical equipment in the photoinhibition experiment. This work was supported by Grant-in-Aids for Scientific Research on Innovative Areas (17H06309 to M.M.) and for Scientific Research (A) (19H01037 to M.M.) from the Ministry of Education, Culture, Sports, Science, and Technology, Japan; and by AMED (JP18dm0207027 to M.M.), a JSPS Fellowship (201801565 to M.K.), and JSPS KAKENHI (JP18J01565 to M.K.) from the Japan Society for the Promotion of Science.

