## Supplementary material for "Independent representations of reward-predicting cues and reward history in frontal cortical neurons": Suppl. figre and table and the legend

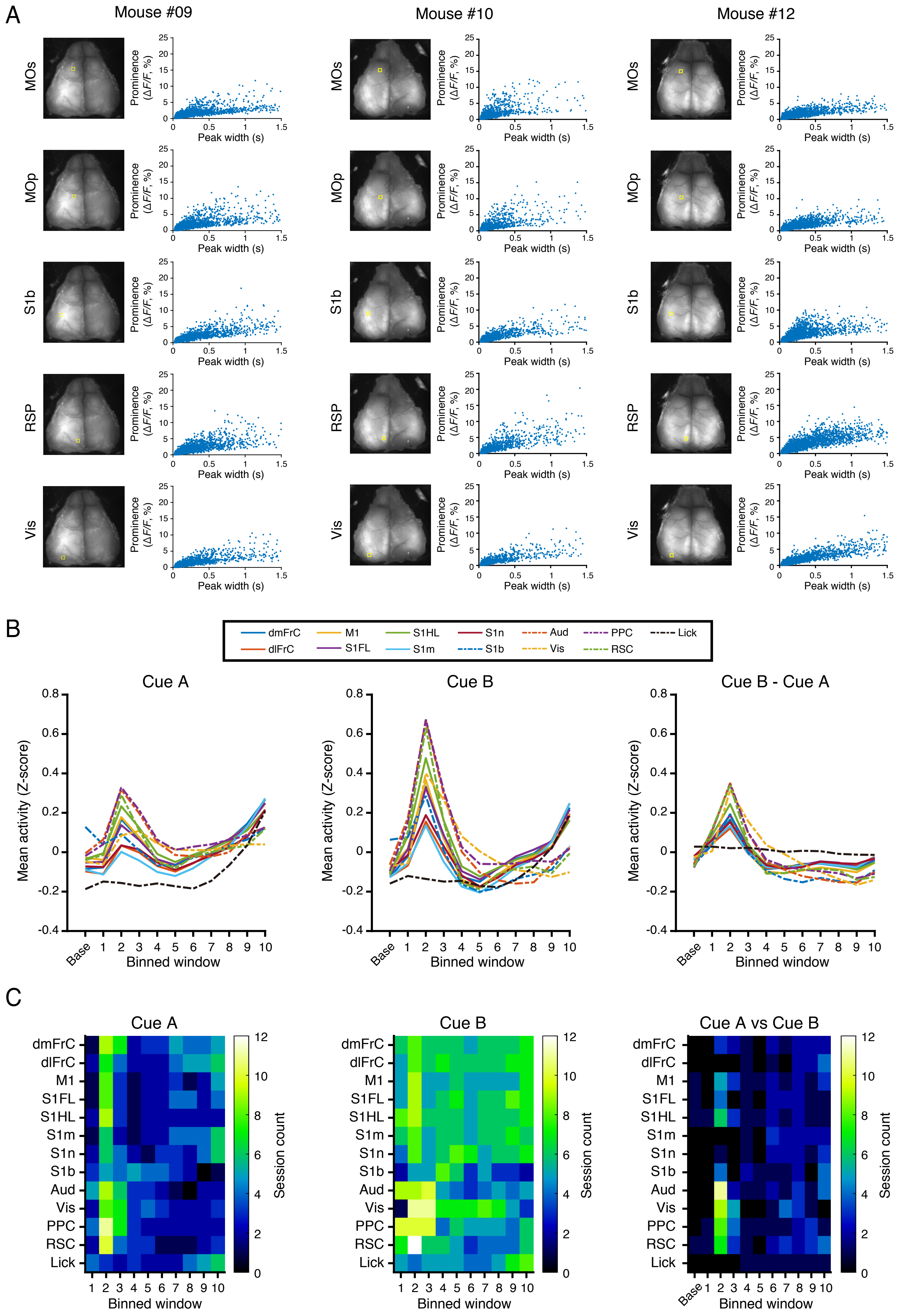
A
Mouse #09
Mouse #10
Mouse #12
MOs
MOp
S1b
RSP
Vis
Prominence (ΔF/F, %)
Peak width (s)
B
dmFrC
dlFrC
M1
S1FL
S1HL
S1m
S1n
S1b
Aud
Vis
PPC
RSC
Lick
Cue A
Cue B
Cue B - Cue A
Mean activity (Z-score)
Binned window
Base
C
Cue A
Cue B
Cue A vs Cue B
Session count
Binned window

**Figure S1, related to the experimental procedure, Figures 1 and 2. Absence of abnormal cortical activity in the mice with pan-neuronal Cre-driven TITL-R-CaMP1.07 and the neuronal response of each dorsal cortical area to two tone cues in pretraining sessions**

(A) Examples of scattered plots of $\Delta F/F$ against peak width in the frontal cortex, primary motor cortex (M1), barrel cortex (S1b), retrosplenial cortex (RSC), and visual cortex (Vis). The values were calculated from spontaneous activities in four Emx1-Cre::CaMKII-tTA::TITL-R-CaMP1.07 mice. Abnormally narrow neurons with large-amplitude activity have been detected in Emx1::CaMKII-tTA::TIGRE-GCaMP6f mice (Ai93 mice crossed with CaMKII-tTA mice and Emx1-Cre mice), with these neurons being assumed to reflect epileptic cortical activity (Steinmetz et al., 2017). However, such neurons were not observed in the transgenic mice that we used.

(B) Normalized neuronal activity of each area and the lick-rate in the pretraining sessions. The neuronal activities and lick-rate during 2 s cue and delay periods were averaged for each 200 ms-bin window. The baseline (base) activity and lick-rate were those averaged over the 500 ms before the cue onset.

Left, cue A trials; middle, cue B; right, the activity after cue A onset subtracted from that after cue B onset.

(C) Left and middle, the number of sessions showing significant differences ($P < 0.01$ with FDR correction) in lick-rate or neuronal activity in each area between the baseline window and each time-bin (total = 12; four mice × three sessions) in cue A (left) and cue B (middle) trials. Right, the number of sessions that showed large differences in neuronal activity between cue A and B trials in each area for each time-bin. The right panels in (B, C) indicate that the sensory response to cue B was higher than that to cue A.

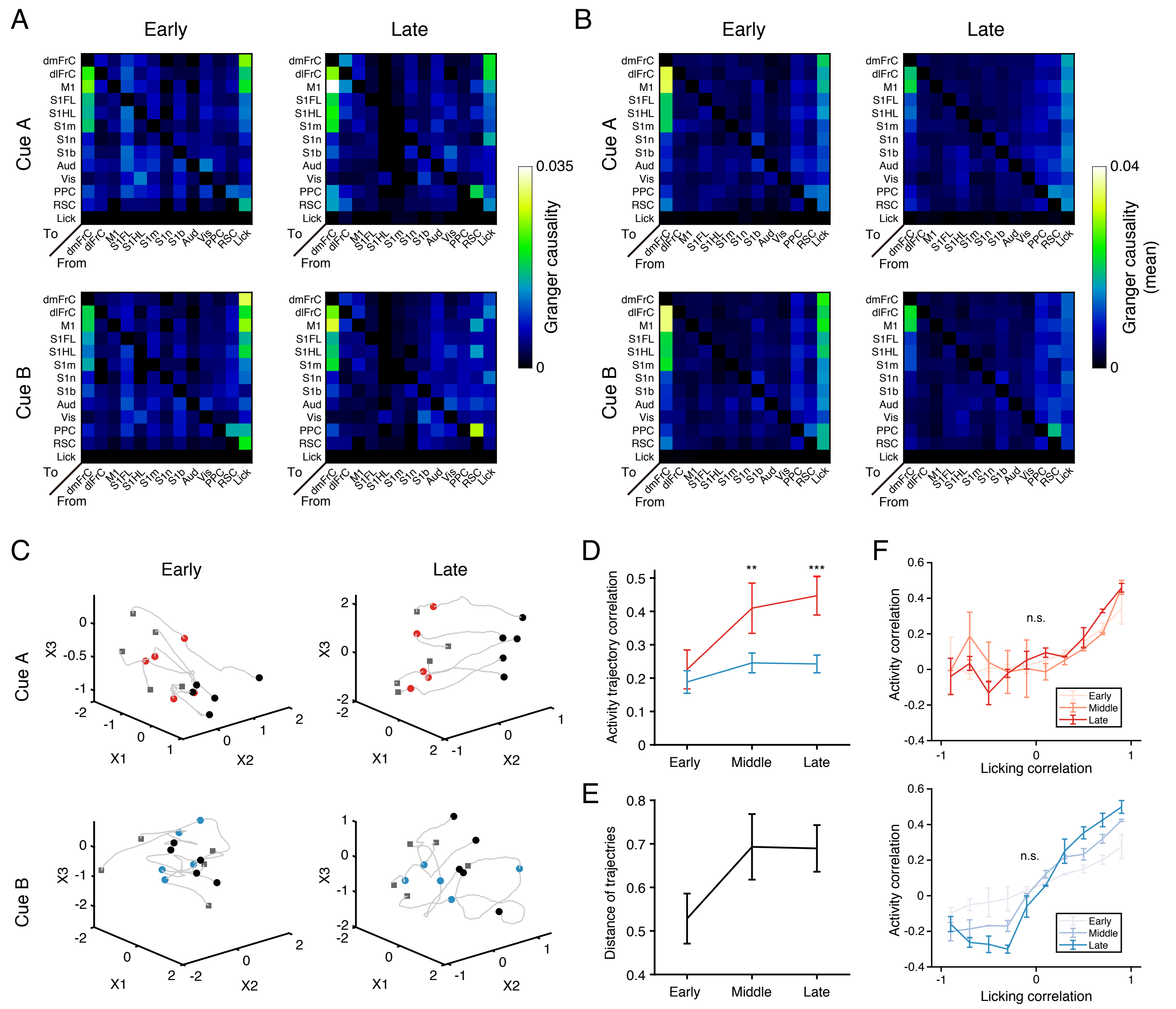


**Figure S2, related to Figure 3. An alternative illustration of Granger causality and the trial-by-trial variability of the cortex-wide neuronal trajectory and licking behavior**

(A, B) Granger causality matrix consisting of the neuronal activity in the twelve areas and lick-rate. The direction of causality is from the area on the horizontal axis to the area on the vertical axis. An example from one animal (A) and the average over all animals (B; the same data used in Figure 3C). Thresholding was not performed.

(C) Representative neuronal trajectories in three-dimensional state space obtained with the factor analysis. The neuronal activities of 12 areas during the 2 s period after the cue onset were decomposed with five factors, and the trajectories of five randomly chosen trials are displayed in the space composed of the first three factors (X1 to X3). Square symbols indicate the time points 1 s before the cue onset. Red and blue circles are the cue A and cue B onset timings, respectively. Black circles are those 2 s after the cue onset.

(D) Correlation of the neuronal trajectories between trials in each cue in early, middle (sessions 4 and 5), and late stages of learning. Red, cue A. Blue, cue B. **: $P < 0.01$, ***: $P < 0.0001$, Wilcoxon rank-sum test with FDR correction.

(E) Distance of the neuronal trajectories between cue A and cue B trials in the early, middle, and late stages of learning.

(F) Dependency of the trial-by-trial correlation in the neuronal trajectories on the trial-by-trial correlation in the lick-rate in the three stages of learning. Top, cue A trials. Bottom, cue B trials. The relationship between the neuronal trajectory and lick-rate was not significant in any learning stage in either cue trial (Friedman's test).

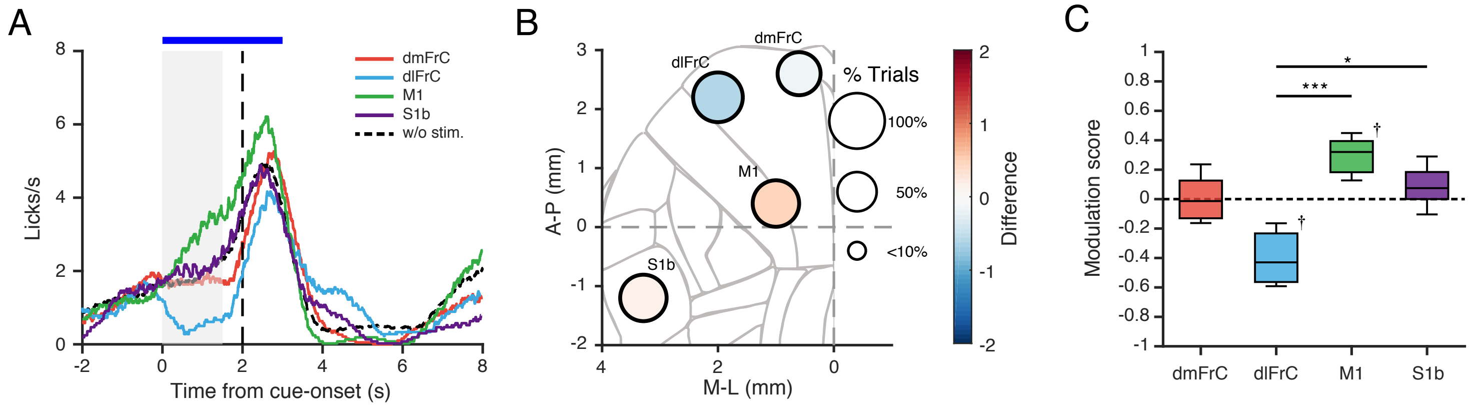


**Figure S3, related to Figure 5. Photoinhibition of neuronal activities in pretraining sessions**

(A) Representative example of the lick-rate in one session. Blue light was bilaterally illuminated on each of the four areas with a probability of 50%. The blue horizontal bar indicates the 3 s photoinhibition period. Red: dmFrC stimulation, blue: dlFrC stimulation, green: M1 stimulation, purple: vS1 stimulation, black dashed line: no stimulation. Each trace is trial-averaged. The SEMs of each trace are not indicated in the visualization.

(B) Representative photoinhibition-induced changes in lick-rate during the cue presentation and delay periods. The color of the circle indicates the difference in lick-rate between the presence and absence of photoinhibition. The size of the circle corresponds to the fraction of trials showing increases or decreases over the mean lick-rate without stimulation (depends on the color of the circle) for all photoinhibited trials.

(C) Photoinhibition-induced modification of lick-rate. The modulation score is a positive or negative value if the lick-rate increased or decreased, respectively. †: $P < 0.05$, comparison between the scores from zero in each photoinhibition site, sign test with FDR correction. *: $P < 0.05$, ***: $P < 0.0001$, Friedman test with multiple correction by dunn-sidak method.

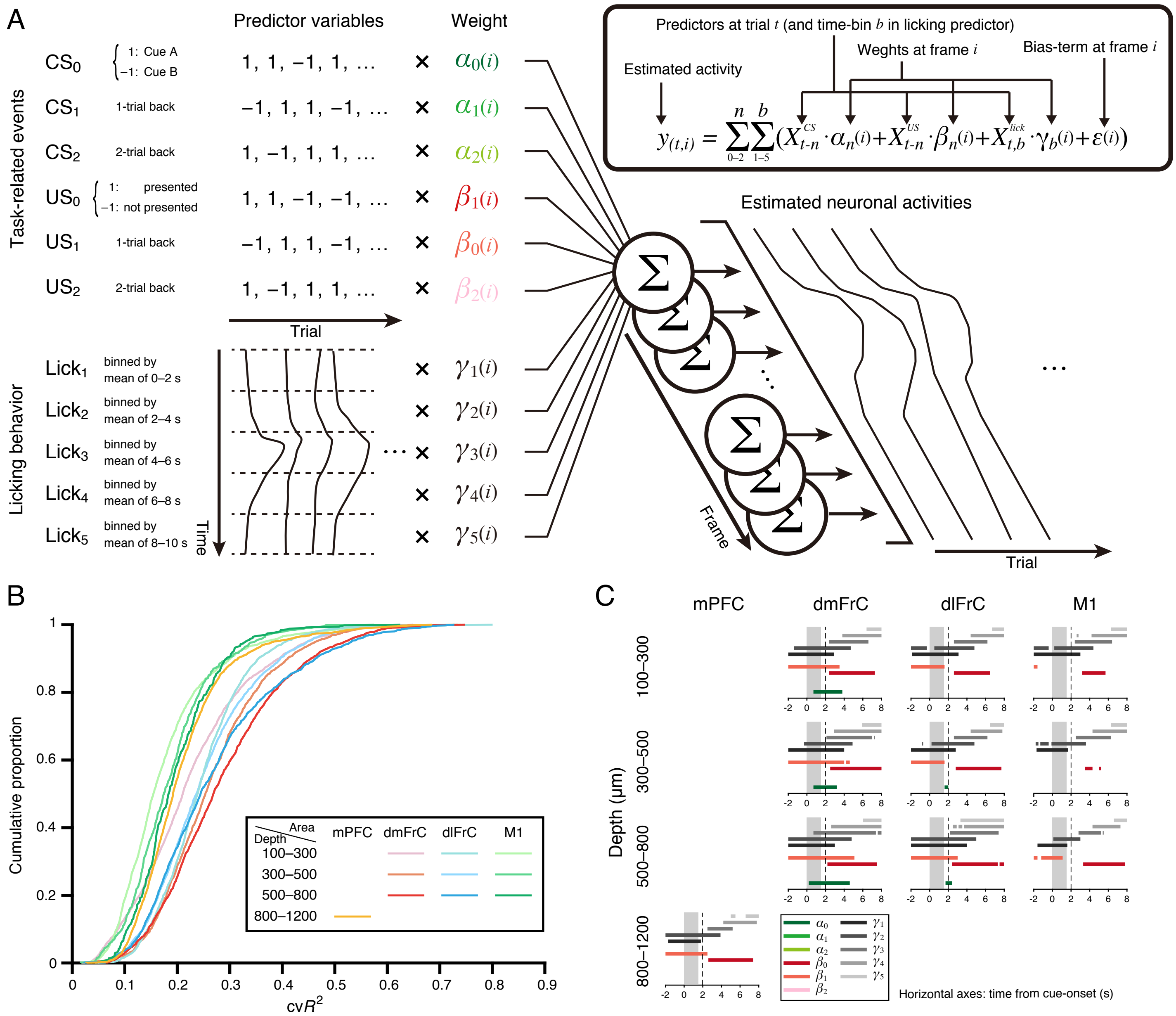


$$y_{(t,i)} = \sum_{0\text{–}2}^{n}\sum_{1\text{–}5}^{b}\left(X^{CS}_{t\text{-}n}\cdot\alpha_n(i)+X^{US}_{t\text{-}n}\cdot\beta_n(i)+X^{lick}_{t,b}\cdot\gamma_b(i)+\varepsilon(i)\right)$$

**Figure S4, related to the experimental procedure and Figure 7. Additional details of the task-encoding model with multiple linear regression**

(A) Schematics of the encoding model with multiple linear regression. CS type (1: cue A, –1: cue B) and US presence (1: presence of reward, –1: absence of reward) were used as task-related predictors. To consider the history of these parameters, the CS type and US presence in one-trial-back and two-trials-back were also included in the model. To consider the influence of licking behavior, the lick-rate in each trial was divided every 2 s, and five bin-averaged lick-rates were used as movement predictors. The activity in each neuron was regressed by estimating a weight vector for each frame and finally concatenating the frame-to-frame estimations to full-frame time-series (upper right inset).

(B) Prediction accuracy of the encoding model estimated with 5-fold cross-validation. The accuracy obtained by squaring the linear correlation between the actual value of the test data and the response of the estimated model was used to indicate goodness of fit ($cvR^2$). The cumulative probability was plotted for each field and each depth. Red, blue, green, and yellow lines indicate dmFrC, dlFrC, M1, and mPFC, respectively. The color depth indicates the cortical depth, with pale colors corresponding to shallow layers and deep colors to deep layers.

(C) The bold lines at each cortical depth in each field indicate the times at which the proportion of neurons with large coefficients of the predictors were above chance level ($P < 0.01$, binomial test).

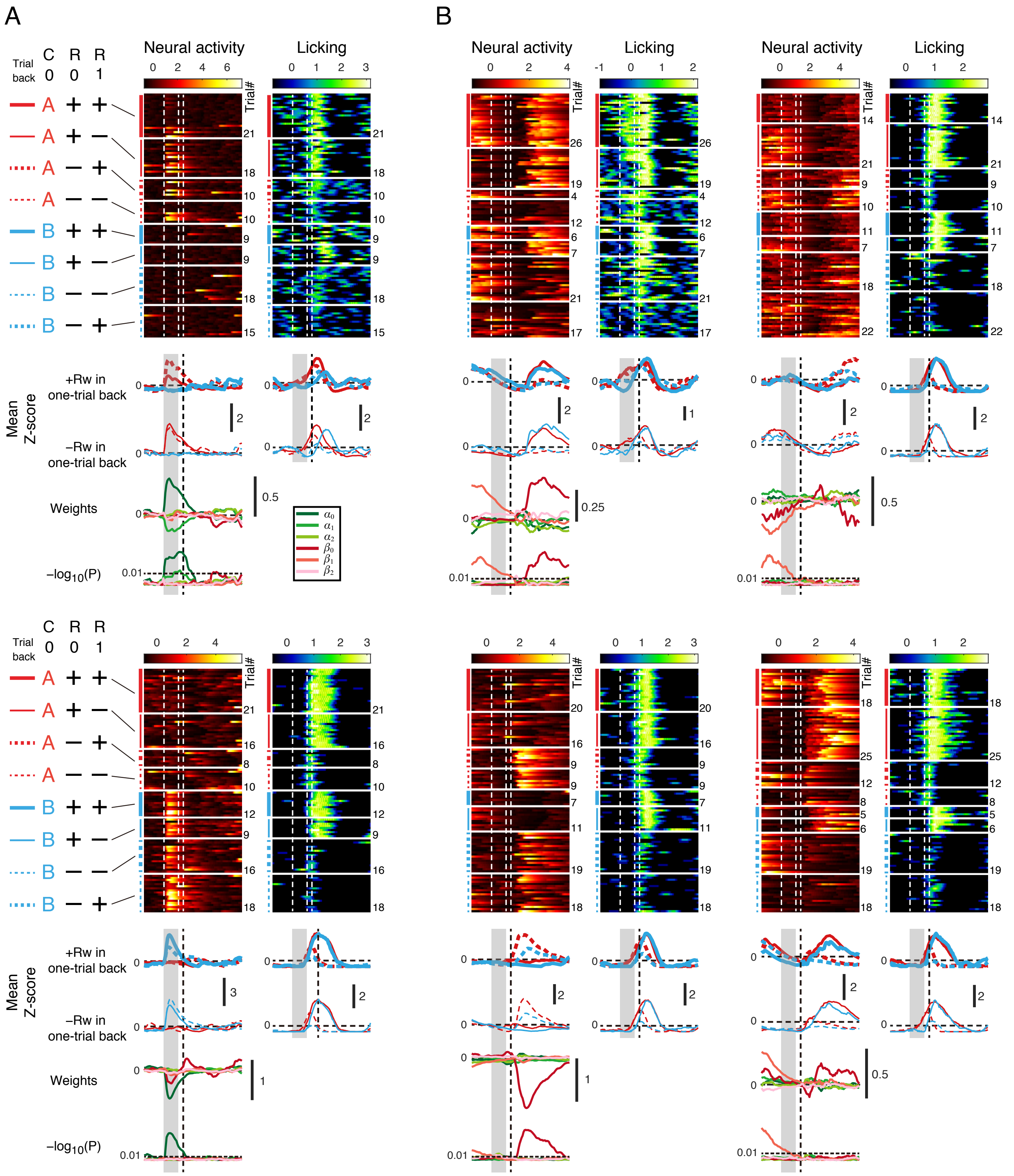


**Figure S5, related to Figure 6. Examples of cue- and reward-encoding neurons that were not licking-related, as revealed by the task-encoding model**

(A, B) Examples of six neurons that were not licking-related but that had task-related activity. Two cue-related neurons (A) and four reward-related neurons (B). Top and bottom panel(s) correspond to neurons that were positively and negatively modulated by cues A and B in (A), or the presence and absence of reward in (B), respectively. The conventions are the same as in Figure 6A.

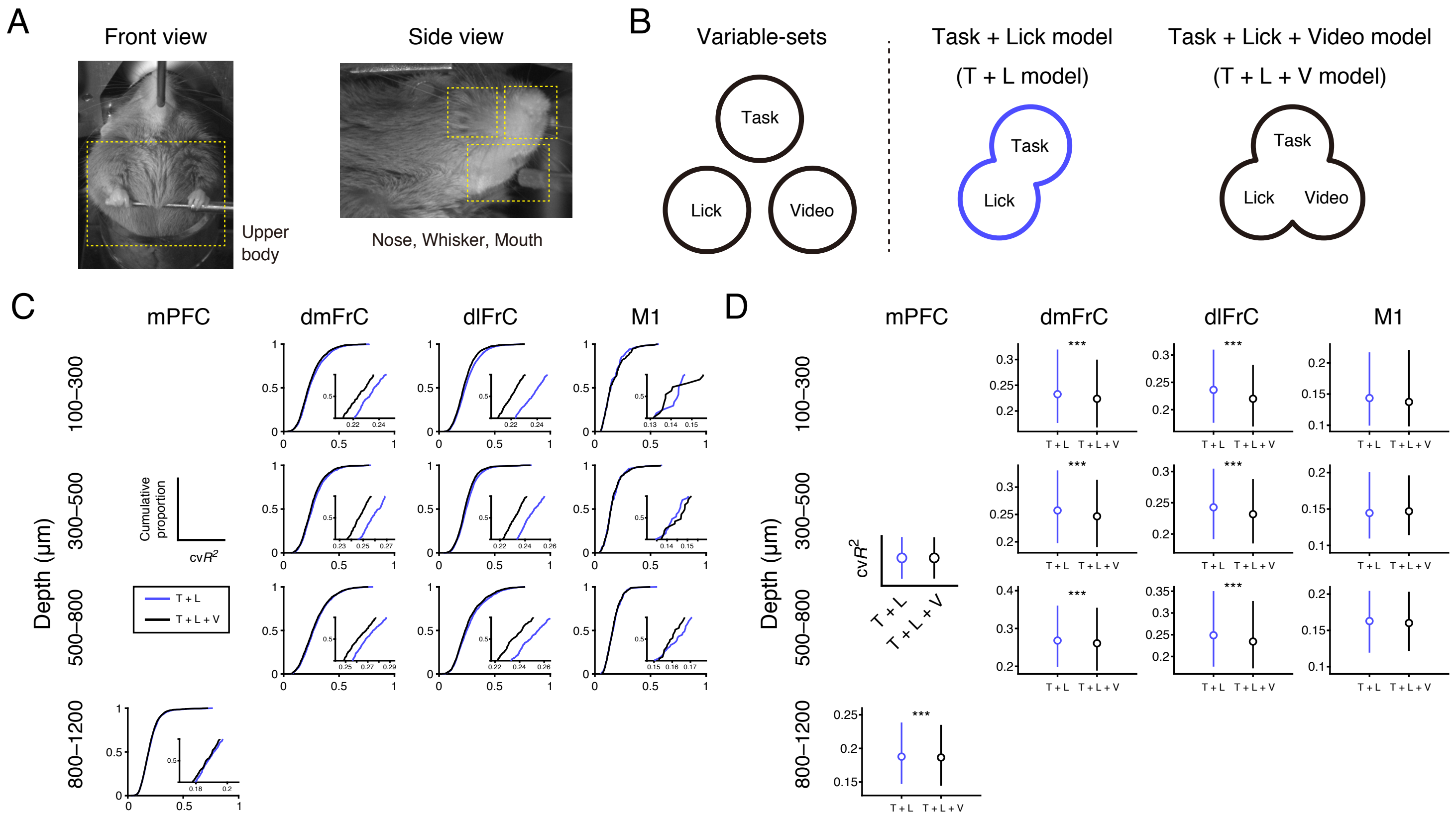
A
Front view
Side view
Upper body
Nose, Whisker, Mouth
B
Variable-sets
Task
Lick
Video
Task + Lick model
(T + L model)
Task + Lick + Video model
(T + L + V model)
C
mPFC
dmFrC
dlFrC
M1
Depth (µm)
100–300
300–500
500–800
800–1200
Cumulative proportion
cvR2
T + L
T + L + V
D
***

**Figure S6, related to Figure 7. The encoding model with task-related and licking-related variables explains the neural activity at an approximately equivalent level to the full model also including body movement-related variables**

(A) Monitoring of the body movements during the task. Upper body and face movements were monitored with two high-speed video cameras. Four ROIs (yellow dotted boxes; upper body, nose, whiskers, and mouth) were set to calculate the movements of specific body parts as body movements. The body movements were calculated as motion-energy values, which were absolute values of the frame-to-frame differences in the pixel intensities of a raw movie, and then the movements in each ROI were decomposed by an SVD-based algorithm and the decomposed values were used for the regression analysis.

(B) Variable-sets contained in the full model and the combinations for partial and full models. Left: the Task variable-set (cue- and reward outcome-histories, $2 \times 3 = 6$ variables), Lick variable-set (z-scored lick-rate segmented and averaged for every 2 s interval in the current trial, five variables), and Video variable-set (most dominant motion-energy in each video ROI segmented and averaged for every 2 s interval in the current trial, $5 \times 4 = 20$ variables) were used for this analysis. Right: two encoding models with and without the Video variable-set. The neural responses of each cell were regressed with the task-encoding model with the Task and Lick variable-sets (T + L), and the full model with the Task, Lick, and Video variable-sets (T + L + V), to estimate the contribution of the video variable-set. Note that the T + L model was the same configuration as that in the main results.

(C, D) Comparison of the prediction accuracy between the task-encoding model and full model. C, Cumulative proportions of cross-validated $R^2$ (cv$R^2$), an estimation of the power of the model for the actual neural responses of all neurons. cv$R^2$ values were separately presented by the region and depth of each neuron. Insets, magnifications around the median values of cv$R^2$. Cyan and black lines correspond to the proportions of T + L and T + L + V models, respectively. D, Summarized results of $cvR^2$ at the population-level. Circles show the median values over the neuron population in each cortical region and depth. The top and bottom edges of lines correspond to the first and third quartiles. Cyan and black correspond to the data calculated from the T + L and T + L + V models, respectively. ***: $P < 0.001$, Wilcoxon sign-rank test with FDR corection. n = 1862 cells in mPFC, 1653 at the depth of 100–300 µm in dmFrC, 1435 at the depth of 300–500 µm in dmFrC, 1856 at the depth of 500–800 µm in dmFrC, 1928 at the depth of 100–300 µm in dlFrC, 2669 at the depth of 300–500 µm in dlFrC, 1484 at the depth of 500–800 µm in dlFrC, 117 at the depth of 100–300 µm in M1, 183 at the depth of 300–500 µm in M1, and 255 at the depth of 500–800 µm in M1. These data were obtained from the all sessions with the behavioral video recording.

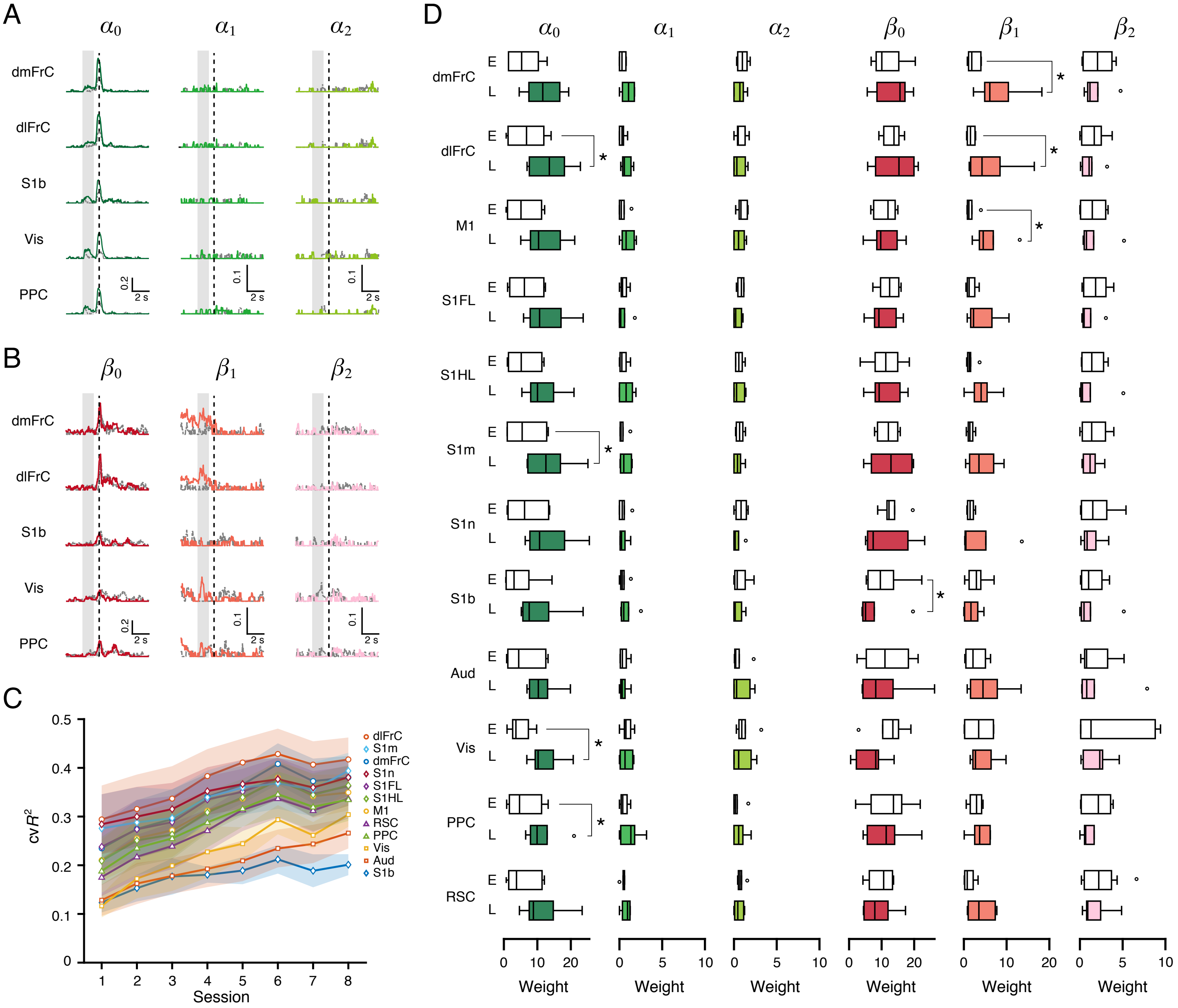

A
$\alpha_0$
$\alpha_1$
$\alpha_2$
dmFrC
dlFrC
S1b
Vis
PPC
0.2
2 s
0.1
2 s
0.1
2 s
B
$\beta_0$
$\beta_1$
$\beta_2$
dmFrC
dlFrC
S1b
Vis
PPC
0.2
2 s
0.1
2 s
0.1
2 s
C
cv$R^2$
0.5
0.4
0.3
0.2
0.1
0
1
2
3
4
5
6
7
8
Session
dlFrC
S1m
dmFrC
S1n
S1FL
S1HL
M1
RSC
PPC
Vis
Aud
S1b
D
$\alpha_0$
$\alpha_1$
$\alpha_2$
$\beta_0$
$\beta_1$
$\beta_2$
dmFrC
dlFrC
M1
S1FL
S1HL
S1m
S1n
S1b
Aud
Vis
PPC
RSC
E
L
*
0
10
20
0
5
10
Weight

**Figure S7, related to Figures 2 and 7. Task-encoding model for cortex-wide neuronal activities and the performance during learning, and changes in the regression weights of cue and outcome variables over different sessions in the wide-field imaging data**

(A, B) Time courses of the mean weights (cue weights $\alpha_0$–$\alpha_2$, A; reward weights $\beta_0$– $\beta_2$, B) for dmFrC, dlFrC, S1b, Vis, and PPC calculated from the wide-field calcium imaging data of six animals. Gray lines indicate the weights in the early stage of learning, and other color lines indicate those in the late stage of learning. Gray shading indicates the cue presentation period. To facilitate clear data visualization, the SEM of each trace is not indicated.

(C) Changes in the prediction accuracy of the model over eight sessions in each cortical area. The neuronal activity for each session was regressed by the task-encoding model, and the prediction accuracy (the square of the linear correlation coefficient between the model response to the test data and the actual response) was calculated with 5-fold cross-validation (cv$R^2$). The data shown are the mean ± SEM from six animals.

(D) The sum of absolute values of each weight in all time frames. The columns correspond to each task-related regressor. The open boxes are the results in the early stage of learning (E), and the colored boxes are the results in the late stage (L). The edges of boxes indicate the first and third quartiles, the inner line is the median over six mice, the whiskers represent minimum and maximum values except for outliers, and the white circles represent outliers outside 1.5 times the IQR. *: $P < 0.05$, Wilcoxon signed-rank test with FDR correction.

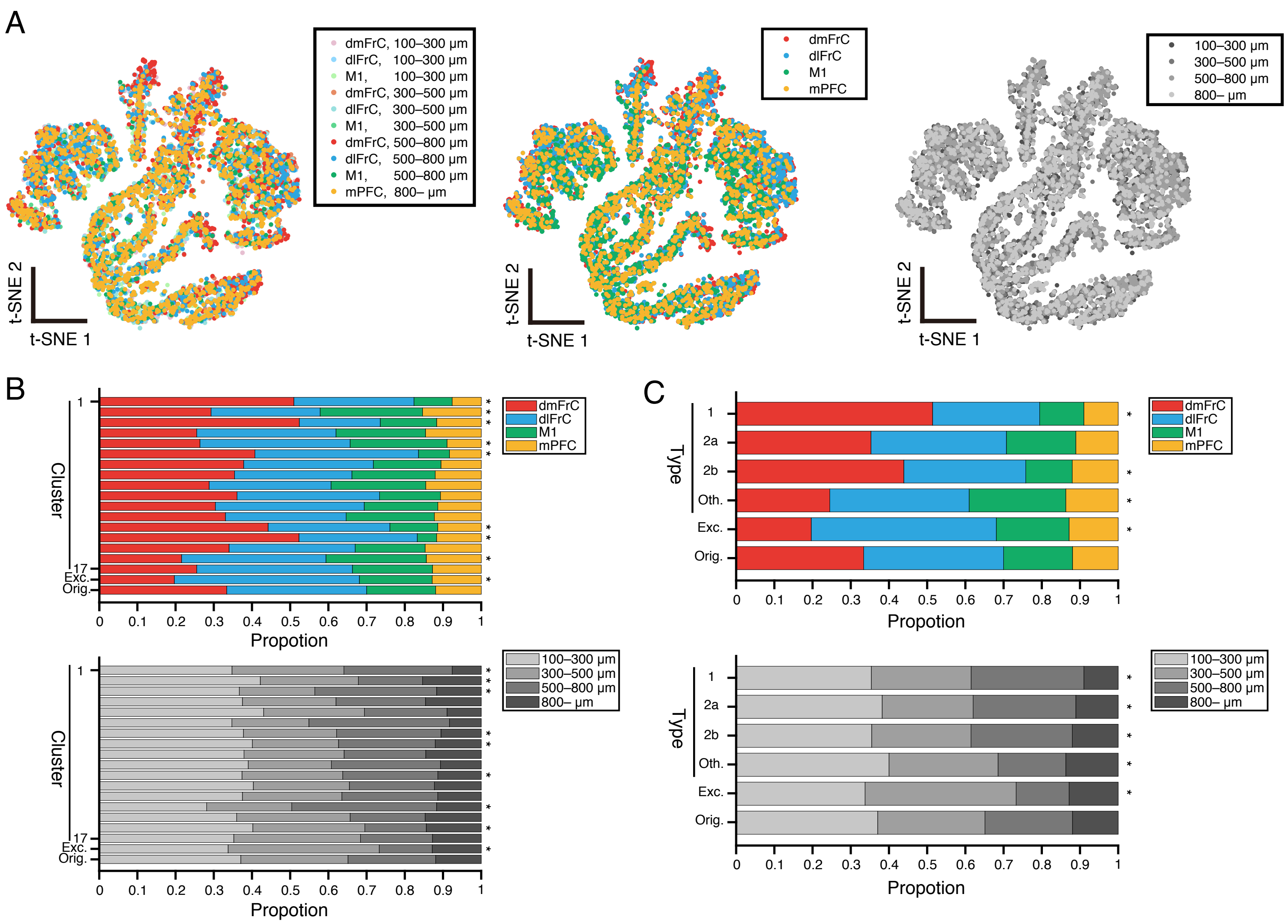


**Figure S8, related to Figure 8. Imaging fields and depths of neurons clustered according to encoding modality**

(A) Each neuron on the t-SNE map shown in Figure 8A was labeled with the corresponding fields and depths (left), fields (middle), and depths (right). Any color that does not appear to be unevenly distributed suggests that the neurons at a specific depth in a specific field did not occupy any cluster.

(B) Ratios of the fields (top) and depths (bottom) in each cluster. Each bar shows the ratio in clusters 1 to 17 from the top, the group of excluded neurons (exc.), and all imaged neurons (orig.). Asterisks indicate that the neuron ratio of the four areas was significantly different from the ratio in all imaged neurons at $P < 0.05$ (chi-square test).

(C) Ratios of the fields (top) and depths (bottom) in types 1, 2a, and 2b. Asterisks indicate that the neuron ratio of the four areas was significantly different to the ratio in all imaged neurons at $P < 0.05$ (chi-square test). The proportions of dmFrC neurons in types 1 and 2b were higher than those in all imaged neurons. The proportions of deep-layer (500–800 µm) neurons in types 1, 2a, and 2b were higher than in all imaged neurons.

| Cluster # | Encoding magnitude of predictor variables: $\alpha_0$ | $\alpha_1$ | $\alpha_2$ | $\beta_0$ | $\beta_1$ | $\beta_2$ | Mean total encoding magnitude | Assigned feature type |
|---|---|---|---|---|---|---|---|---|
| 1 | 7.768 | 0.7046 | 0.4813 | 1.019 | 2.136 | 0.7055 | 12.81 | 1 |
| 2 | 1.634 | 0.8826 | 0.6773 | 0.03808 | 0.1838 | 0.3494 | 3.766 | Other |
| 3 | 7.56 | 0.5277 | 0.5253 | 0.622 | 1.666 | 1.067 | 11.97 | 1 |
| 4 | 2.732 | 0.7599 | 0.2991 | 0.07808 | 0.2519 | 0.3884 | 4.51 | Other |
| 5 | 0.9805 | 0.8255 | 0.8774 | 0.04187 | 0.09289 | 0.3335 | 3.152 | Other |
| 6 | 1.071 | 0.4087 | 0.336 | 10.94 | 1.968 | 0.5536 | 15.28 | 2a |
| 7 | 0.8729 | 0.4114 | 0.3915 | 10.46 | 4.096 | 0.6021 | 16.83 | 2a |
| 8 | 0.7811 | 0.4327 | 0.3449 | 6.938 | 1.263 | 0.6138 | 10.37 | 2a |
| 9 | 0.5614 | 0.3819 | 0.3898 | 4.159 | 1.74 | 0.5683 | 7.801 | 2a |
| 10 | 1.225 | 0.5134 | 0.4336 | 10.86 | 1.88 | 0.7308 | 15.65 | 2a |
| 11 | 0.8531 | 0.568 | 0.4659 | 7.084 | 2.201 | 0.7372 | 11.91 | 2a |
| 12 | 0.7095 | 0.4653 | 0.3977 | 5.735 | 1.222 | 0.5993 | 9.129 | 2a |
| 13 | 0.68 | 0.4957 | 0.3332 | 0.6184 | 6.776 | 0.8066 | 9.71 | 2b |
| 14 | 1.099 | 0.5608 | 0.3517 | 0.7471 | 9.945 | 1.22 | 13.92 | 2b |
| 15 | 0.2644 | 0.5212 | 0.3656 | 0.09873 | 2.931 | 1.276 | 5.457 | 2b |
| 16 | 0.05916 | 0.5046 | 0.5911 | 0.0009926 | 0.2141 | 0.6839 | 2.054 | Other |
| 17 | 0.0634 | 0.6694 | 0.5268 | 0.002398 | 0.4933 | 1.899 | 3.654 | Other |

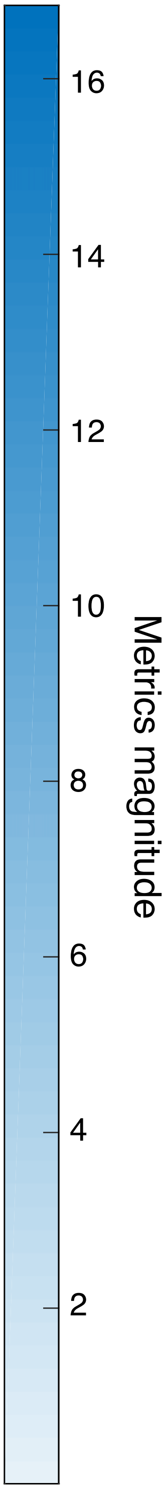


**Table S1. Encoding metrics and the order and assigned group of sorted clusters.**

The clusters were sorted according to encoding magnitude and sign (see Methods). The first six columns from the left show the average encoding magnitude of each cluster. The seventh column corresponds to the neuron-averaged total encoding magnitude, calculated by temporal summation of the six predictor variables (see Methods). The eighth column indicates the assigned cluster type.

**Video S1. Representative two-photon time-lapse imaging planes in four frontal cortical regions.**

Examples of functional imaging in the four frontal cortices expressing jRGECO1a during the classical conditioning task. Top left, top right, bottom left, and bottom right movies correspond to time-lapse movies taken from mPFC, dmFrC, dlFrC, and M1, with depths from the cortical surface of 980, 770, 430, and 150 μm, respectively. The field of view is 509.12 × 509.12 μm (512 × 512 pixels). The circle in each panel indicates the cue presentation and the red or blue color corresponds to the cue A or B presentation, respectively. The white diamond in each panel indicates the timing of water delivery. The original data were acquired at a 30 Hz frame rate, with the movies being down-sampled to 10 Hz by three-frame averaging.
